# Conformational Landscaping and Dynamic Mutational Profiling of Binding Interactions and Immune Escape for Broadly Neutralizing Class I Antibodies with SARS-CoV-2 Spike Protein: Distributed Binding Hotspot Networks Underlie Mechanism of Viral Resistance Against Existing Variants

**DOI:** 10.1101/2025.06.07.658468

**Authors:** Mohammed Alshahrani, Vedant Parikh, Brandon Foley, Gennady Verkhivker

## Abstract

The rapid evolution of SARS-CoV-2 has underscored the need for a detailed understanding of antibody binding mechanisms to combat immune evasion by emerging variants. In this study, we investigated the interactions between Class I neutralizing antibodies BD55-1205, BD-604, OMI-42, P5S-1H1, and P5S-2B10 and the receptor-binding domain (RBD) of the SARS-CoV-2 spike protein using multiscale modeling which combined coarse-grained simulations and atomistic reconstruction of conformational landscapes together with mutational scanning of the binding interfaces, dynamic profiling of binding and immune escape using molecular mechanics generalized Born surface area (MM-GBSA) analysis. A central theme emerging from this work is the critical role of epitope breadth and interaction diversity in determining an antibody resilience to mutations. BD55-1205 antibody exemplifies the advantages of broad epitope coverage and distributed hotspot mechanisms. By engaging an extensive network of residues across the RBD, BD55-1205 minimizes its dependence on individual side-chain conformations, allowing it to maintain robust binding even when key residues are mutated. This adaptability is particularly evident in its tolerance to mutations at positions such as L455 and F456, which severely compromise other antibodies. The ability of BD55-1205 to sustain cumulative interactions underscores the importance of targeting diverse epitopes through multiple interaction mechanisms, a strategy that enhances resistance to immune evasion while maintaining functional integrity. In contrast, BD-604 and OMI-42, with localized binding mechanisms, are more vulnerable to escape mutations at critical positions such as L455, F456, and A475. P5S-1H1 and P5S-2B10 exhibit intermediate behavior, balancing specificity and adaptability but lacking the robustness of BD55-1205. Mutational scanning identified key residues Y421, Y489, and F456 as critical hotspots for RBD stability and antibody binding, highlighting their dual role in viral fitness and immune evasion. The computational predictions generated through mutational scanning and MM-GBSA analysis demonstrate excellent agreement with experimental data on average antibody escape scores. This study underscores the diversity of binding mechanisms employed by different antibodies and molecular basis for high affinity and excellent neutralization activity of the latest generation of antibodies.

## Introduction

Structural and biochemical studies of the SARS-CoV-2 Spike (S) glycoprotein have unveiled critical insights into the mechanisms driving viral transmission, immune evasion, and host-cell entry [1–9]. The S glycoprotein is distinguished by its remarkable conformational flexibility, particularly within the S1 subunit, which encompasses several key domains: the N-terminal domain (NTD), the receptor-binding domain (RBD), and two structurally conserved subdomains, SD1 and SD2. This intrinsic flexibility enables the S glycoprotein to dynamically adapt to different stages of the viral entry process, enhancing both its functionality and ability to evade immune detection [1–9]. The NTD facilitates the initial attachment to host cells, while the RBD plays a pivotal role in binding to the angiotensin-converting enzyme 2 (ACE2) receptor, a crucial step for viral entry [10–15]. Simultaneously, the SD1 and SD2 subdomains stabilize the prefusion conformation of the S glycoprotein and assist in its transition to the postfusion state, a necessary process for membrane fusion and subsequent infection [10–15]. Conformational changes in the NTD and RBD drive transitions between the closed and open states of the S protein, allowing the virus to efficiently engage with host receptors while leveraging structural variability to evade immune surveillance. This dynamic behavior not only enhances the virus infectivity but also contributes to its high transmissibility and pathogenicity by enabling immune evasion [10–15]. Biophysical investigations have further clarified the thermodynamic and kinetic principles governing the functional transitions of the S protein [[16–18]]. These studies reveal that mutations within the S protein, particularly in the S1 subunit, can induce structural alterations that affect its stability and conformational dynamics. Such changes critically influence the protein ability to toggle between open and closed states, thereby modulating the accessibility of the RBD—a key determinant for viral attachment to host cells. Moreover, the structural variability introduced by these mutations enhances the virus capacity to evade immune responses, complicating the host ability to mount an effective defense [16–18]. Extensive cryo-electron microscopy (cryo-EM) and X-ray structures of SARS-CoV-2 S protein variants of concern (VOCs) in various functional states, along with their interactions with antibodies, highlight how VOCs can induce structural changes in the dynamic equilibrium of the S protein [19–25]. These findings underscore the balance between structural stability, immune evasion, and receptor binding that shapes the evolutionary trajectory of SARS-CoV-2 and its variants.

The emergence and evolution of SARS-CoV-2 variants such as XBB.1 and XBB.1.5 have garnered significant attention due to their enhanced growth advantages, transmissibility, and immune evasion capabilities. These subvariants represented critical milestones in the virus evolutionary trajectory, highlighting its ability to adapt through mutations that optimize receptor binding while evading immune defenses [26–28]. XBB.1.5, a descendant of the BA.2 lineage, arose through recombination events that introduced key mutations in the RBD of the S protein. These mutations significantly enhance its binding affinity for the ACE2 receptor, making it more infectious than earlier Omicron strains [26–28]. XBB descendants like EG.5 and EG.5.1 carry an additional F456L mutation contributing to increased immune escape [29–32]. A particularly striking example of convergent evolution is observed in “FLip” variants, which carry both L455F and F456L mutations. These include subvariants such as JG.3, JF.1, GK.3, and JD.1.1, which emerged independently across different lineages [33]. The recurrence of the L455F/F456L double mutation underscores its selective advantage, enabling XBB variants to outcompete other strains within the human population by balancing immune evasion with receptor binding efficiency [33]. Another notable variant, BA.2.86, evolved from the BA.2 lineage and demonstrated significant genetic divergence from earlier forms [34–38]. This variant exhibited heightened immune evasion against RBD-targeted antibodies, surpassing even XBB.1.5 and EG.5.1 in its ability to resist neutralization [34–38]. Its descendant, JN.1, acquired an additional L455S mutation, further enhancing its capacity to evade immune responses. Biochemical studies using surface plasmon resonance (SPR) assays revealed that JN.1 exhibits reduced ACE2 binding affinity but gains a marked advantage in evading class-1 RBD antibodies S2K146 and Omi-18, as well as the class-3 antibody S309 [39]. Subsequent research has identified additional mutations at key positions such as L455, F456, and R346, further illustrating the virus adaptive potential. The “SLip” variant, derived from JN.1, carries the L455S mutation alongside F456L, enabling it to escape neutralizing antibodies more effectively [40–43]. More recently, the “FLiRT” variant has emerged, featuring an additional R346T mutation on the SLip backbone. This mutation enhances the variant ability to evade immune surveillance while maintaining sufficient ACE2 binding affinity [40–43].

The JN.1 subvariants KP.2 and KP.3 have independently acquired key mutations in the S protein, including R346T, F456L, Q493E, and V1104L, which enhance their transmissibility and immune evasion capabilities . Notably, KP.3, a descendant of JN.1 referred to as “FLuQE,” carries additional mutations such as L455S and demonstrates strong growth advantages [44] . Other JN.1 subvariants, including LB.1 and KP.2.3, have also emerged, sharing mutations like S:R346T and S:F456L while acquiring unique changes such as S:S31-and S:Q183H (LB.1) or S:H146Q (KP.2.3). These convergent mutations contributed to increased immune evasion and a higher effective reproduction number [45]. Among these variants, the F456L mutation plays a critical role in enhancing antibody evasion, with KP.3 being one of the most immune-evasive JN.1 sublineage [46]. These findings underscore the rapid evolutionary dynamics of SARS-CoV-2 and its remarkable capacity to adapt through mutations that balance immune evasion with functional constraints. The convergence of mutations like L455F, F456L, and R346T highlights the intense selective pressure driving viral evolution, as the virus continually seeks to optimize its fitness in the face of immune challenges Recent cryo-EM studies of JN.1, KP.2, and KP.3 RBD complexes revealed that F456L mutation enhances the binding potential of Q493E, leading to stronger interactions of KP.3 with the ACE2 receptor [47] . This synergy provides an additional evolutionary advantage, enabling the virus to incorporate additional immune-evasive mutations while maintaining high infectivity.

Meanwhile, XEC, a recombinant variant derived from KP.3 has gained attention due to two additional mutations—F59S and T22N—in the NTD [48–50]. Functional assays indicate that XEC exhibits significantly higher infectivity compared to KP.3 and demonstrates enhanced resistance to immune responses [51,52]. KP.3.1.1 subvariant was derived from KP.3 through deletion at S31 position. Structural analysis of the KP.3.1.1 RBD/ACE2 complex and binding data are consistent with an epistatic interaction between the F456L and Q493E mutations in the RBD of KP.3.1.1, where Q493E alone substantially reduces binding but, in the presence of F456L, these residues synergize to restore ACE2 binding affinity [53,54]. The convergent emergence of mutations F456L and Q493E across JN.1 subvariants highlighted the virus ability to balance immune evasion with receptor binding affinity. Functional studies indicated that KP.3.1.1 exhibits reduced binding affinity to monoclonal antibodies across multiple classes, partly due to S31Δ-introduced N30 glycosylation [53,54]. S31del of KP.3.1.1 and T22N of XEC, which could introduce new N-linked glycans on the NTD. Interestingly, both variants demonstrated increased resistance against monoclonal neutralizing antibodies targeting various epitopes on the RBD, suggesting that the additional NTD glycosylation of KP.3.1.1 and XEC could enhance immune evasion via allosteric effects [49].

The SARS-CoV-2 variant LP.8 carries several notable mutations, including S31del, F186L, Q493E, and H445R, which contribute to its unique evolutionary trajectory. Nextstrain an open-source project for real time tracking of evolving pathogen populations (https://nextstrain.org/). Nextstrain provides dynamic and interactive visualizations of the phylogenetic tree of SARS-CoV-2, allowing users to explore the evolutionary relationships between different lineages and variants. This approach assigns SARS-CoV-2 variant as clade when it reaches a frequency of 20% globally at any time point. A new clade should be at least 2 mutations away from its parent major clade. According to the Nextstrain-based evolutionary analysis (Figure 1, Supplementary Materials, Figure S1-S3, Table S1) a descendant of LP.8, LP.8.1 (Clade 25A/Pango lineage) has undergone further mutations, losing the S31 residue to become a “DeFLiRT” variant—a FLiRT variant that evolved to lose S31 —and subsequently acquiring Q493E and R190S [55]. While XEC and KP.3.1.1 have rapidly outpaced earlier variants like KP.3 to become globally dominant due to their distinctive N-terminal domain (NTD) mutations [53,54], several emerging sublineages of JN.1, such as LF.7.2.1, MC.10.1, NP.1, and LP.8.1, have demonstrated even greater growth advantages [55]. Among these, LF.7.2.1 stands out for its additional A475V mutation compared to its predecessor LF.7, which already harbors S31P, K182R, R190S and K444R mutations (Figure1, Table S1). Similarly, LP.8.1, which features an additional R190S mutation, has surged exhibiting the highest growth advantage among currently circulating variants (Figure 1, Supplementary Materials, Figure S1-S3).

**Figure 1.**
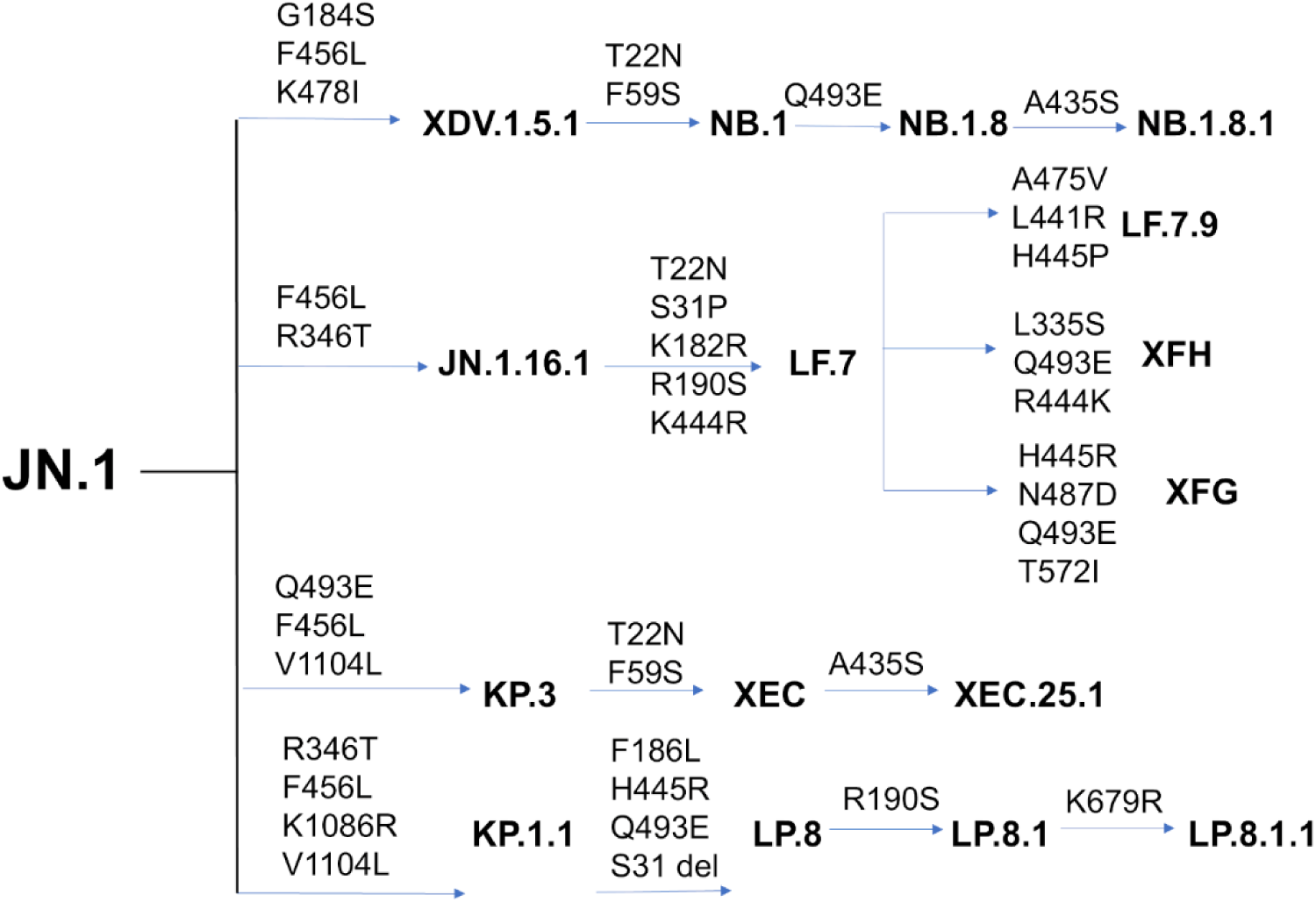
SARS-CoV-2 variant evolution from JN.1, with the main branches of Omicron variants and their respective mutations

Recent studies have elucidated the mechanisms behind the rapid spread of these variants. For instance, LF.7.2.1 demonstrates significantly increased immune evasion compared to XEC, primarily due to the A475V mutation, which enables the virus to escape neutralization by Class 1 antibodies [55]. Meanwhile, LP.8.1 exhibits humoral immune evasion comparable to XEC but shows enhanced engagement efficiency with the ACE2 receptor, supporting its exceptional growth dynamics [55]. These findings highlight the delicate balance between immune evasion and ACE2 binding efficiency in the evolutionary trajectory of SARS-CoV-2, emphasizing how the virus continually optimizes these traits to maintain fitness and transmissibility. The most recent study highlighted the emergence of the SARS-CoV-2 variant BA.3.2, which harbors over 50 mutations relative to its ancestral BA.3 lineage, showing antibody evasion, including resistance to Class 1/4 monoclonal antibodies and yet at the reduced ACE2 engagement [56]. Other variants such as NB.1.8.1, LF.7.9, XEC.25.1, XFH, and XFG displayed enhanced growth advantages over LP.8.1.1 [56]. This study demonstrated that XFG and LF.7.9 feature strong immune escape associated with A475V and N487D mutations but with reduced receptor-binding efficiency. In contrast, NB.1.8.1 featuring mutations G184S, F456L, K478I, T22N, F59S, Q493E and A435S Figure 1, Supplementary Materials, Figure S1-S3) retained high ACE2 affinity and immune evasion, supporting its potential for future dominance. Similarly, XEC.25.1, a derivative of XEC, harbors the A435S mutation and also demonstrates a high growth advantage [56].

The fundamental aspect of the immune response to SARS-CoV-2 is the production of antibodies that target various regions of the S protein, which plays a central role in viral entry into host cells. High-throughput yeast display screening and deep mutational scanning (DMS) have enabled accurate prediction of the escape mutation profiles associated with the RBD residues and mapping of the functional epitopes targeted by neutralizing antibodies leading to a comprehensive classification of distinct epitope groups (A–F) [57,58]. The molecular mechanisms underlying the broadly neutralizing antibodies induced by XBB/JN.1 infections was conducted using high-throughput yeast-display-based DMS assays and the escape mutation profiles in screening of a total 2,688 antibodies, including 1,874 isolated from XBB/JN.1 infection cohorts [59]. In this study, Cao and colleagues showed the possibility of accurately predicting SARS-CoV-2 RBD evolution by aggregating high-throughput antibody DMS results and constructing pseudoviruses that carry the predicted mutations as filters to screen for antibodies. Subsequently, this group identified four E1 group antibodies BD55-3546, BD55-3152, BD55-5585, BD55-5549 and BD55-5840 (SA58) antibodies a well as F3 antibodies BD55-4637, BD55-3372, BD55-5483, and BD55-5514 (SA55) [60,61]. In the recent pioneering investigation, Cao and colleagues leveraged DMS to predict viral evolution and to select for monoclonal antibodies neutralizing both existing and prospective variants. A retrospective analysis of 1,103 SARS-CoV-2 wildtype-elicited monoclonal antibodies shows that this method can increase the probability of identifying effective antibodies against the XBB.1.5 strain from 1% to 40% [62]. Through this screening process they identified BD55-1205, the only wild type (WT)-elicited monoclonal antibodies potent to all major variants. BD55-1205 exhibits exceptional neutralization breadth against the latest variants and prospective mutants including XBB, BA.2.86, and JN.1-derived subvariants, and a high barrier to escape [62]. A yeast-display system combined with a machine learning (ML)-guided approach for library design enabled an investigation of a larger number of Ab variants and the identification of a class 1 human antibody VIR-7229 potently neutralizing EG.5, BA.2.86, and JN.1 [63]. The structures of VIR-7229-bound to XBB.1.5 and EG.5 structures showed that the VIR-7229 interactions can accommodate both F456 and L456 in the corresponding genetic backgrounds and tolerate an extraordinary epitope variability exhibiting high barrier for the emergence of resistance, partly attributed to its high binding affinity [63].

Computer simulations have significantly advanced our understanding of the dynamics and functions of the S protein and S complexes with ACE2 and antibodies at the atomic level. Molecular dynamics (MD) simulations and Markov state models (MSM) have systematically characterized the conformational landscapes of XBB.1 and XBB.1.5 Omicron variants and their complexes [64]. Mutational scanning and binding analysis of the Omicron XBB spike variants with ACE2 and a panel of class 1 antibodies provided a quantitative rationale for the experimental evidence [65,66]. We combined AlphaFold2-based atomistic predictions of structures and conformational ensembles of the S complexes with the ACE2 for the most dominant Omicron variants JN.1, KP.1, KP.2 and KP.3 to examine the mechanisms underlying the role of convergent evolution hotspots in balancing ACE2 binding and Ab evasion [67]. Our recent studies suggested a mechanism in which the pattern of specific escape mutants for ultrapotent antibodies may be driven by a complex balance between the impact of mutations on structural stability, binding strength, and long-range communications [68,69]. Moreover, convergent Omicron mutations can display epistatic couplings with the major stability and binding affinity hotspots which may allow for the observed broad antibody resistance [68,69] Computational studies examined mechanisms of broadly neutralizing antibodies : E1 group (BD55-3152, BD55-3546 and BD5-5840/SA58) and F3 group (BD55-3372, BD55-4637 and BD55-5514/SA55) revealing that E1 and F3 groups of antibodies leverage strong hydrophobic interactions with the binding epitope hotspots critical for the spike stability and ACE2 binding, while escape mutations tend to emerge in sites associated with synergistically strong hydrophobic and electrostatic interactions [70]. In the recent modeling study, a comparative analysis of the binding mechanisms and resistance profiles of S309, S304, CYFN1006, and VIR-7229 revealed distinct molecular strategies employed by these antibodies that can be broadly categorized into two paradigms: (1) conservation-driven binding, where antibodies exploit highly conserved residues critical for viral function, and (2) adaptability-driven binding, where antibodies utilize structural flexibility and compensatory interactions to tolerate mutations while maintaining neutralization efficacy [71]. These findings underscore the remarkable ability of the spike evolution to exploit synergistic energetic contributions at critical hotspots, leading to significant reductions in antibody binding upon mutation. Functionally balanced substitutions that optimize tradeoffs between immune evasion, high ACE2 affinity and sufficient conformational adaptability might be a common strategy of the virus evolution and serve as a primary driving force behind the emergence of new Omicron subvariants [72–74]. At the molecular level, the evolution of immune escape hotspots in SARS-CoV-2 is a complex process driven by antigenic drift and convergent evolution. The virus achieves this through a fine-tuned balance of immune evasion and receptor-binding affinity, influenced by both genetic mutations and the diversity of antibody responses.

In this study, we employed coarse-grained (CG) simulations using CABS-flex approach that efficiently combines a high-resolution coarse-grained model and efficient search protocol capable of accurately reproducing all-atom MD simulation trajectories and dynamic profiles of large biomolecules on a long time scale [75–80]. We performed multiple CG-CABS simulations of the S protein complexes with a panel of broadly neutralizing class I antibodies BD55-1205, BD-604, OMI-42, P5S-1H1 and P5S-2B10 (Figure 2) followed by all-atom reconstruction of trajectories to examine how structural plasticity of the RBD regions can be modulated by binding. We determine specific dynamic signatures induced by antibodies.

**Figure 2.**
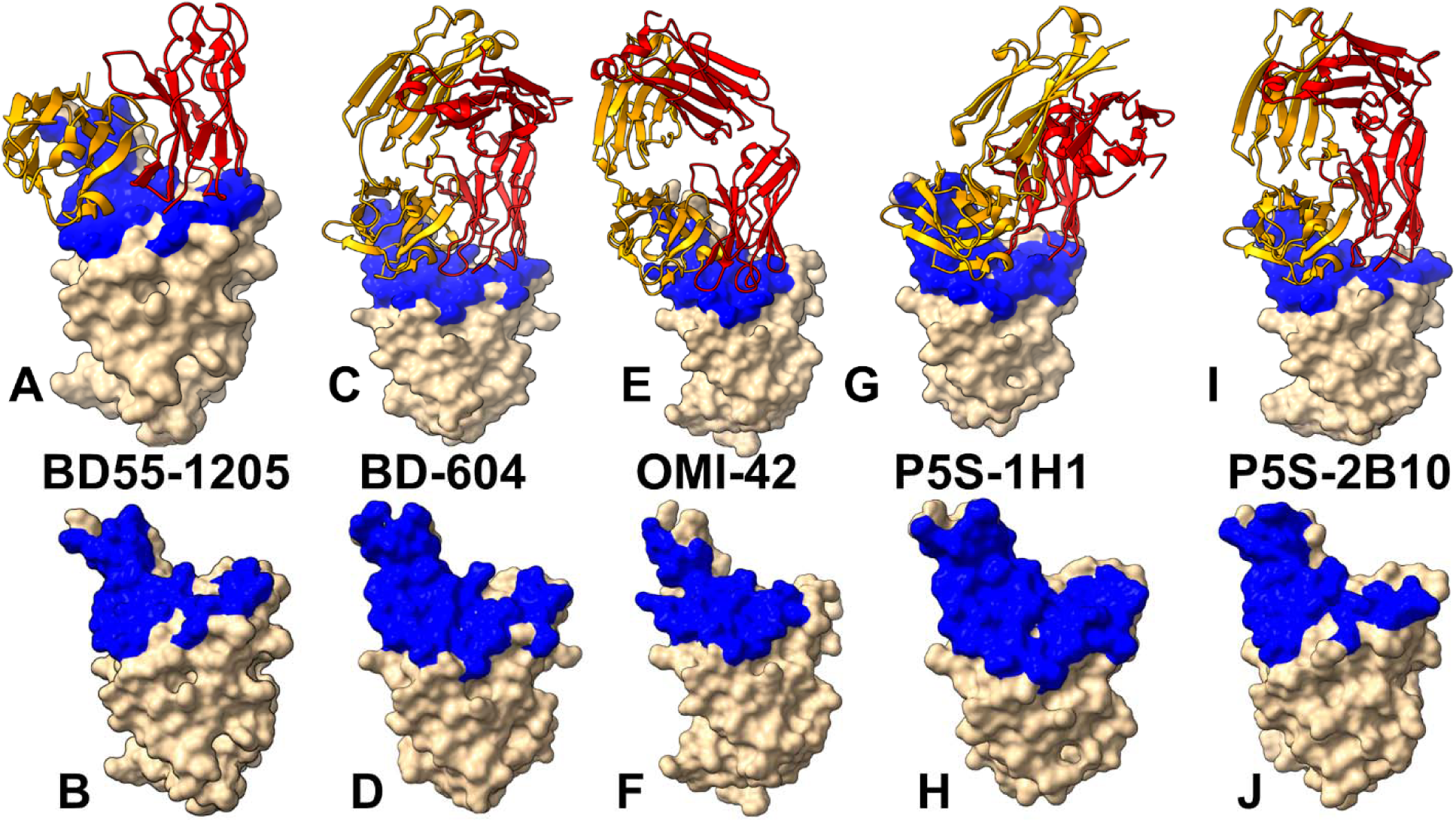
Structural organization of the RBD complexes and binding epitopes for class I antibodies. (A) The structure of BD55-1205 with XBB.1.5 RBD (pdb id 8XE9). The heavy chain in orange ribbons, the light chain in red ribbons. (B) The RBD and binding epitope footprint for BD55-1205. The binding epitope residues are shown in blue surface. (C) The structure of BD-604 bound with RBD (pdb id 7CHF, 7CH4). The heavy chain in orange ribbons, the light chain in red ribbons. (D) The RBD and binding epitope footprint for BD-604. The binding epitope residues are shown in blue surface. (E) The structure of OMI-42 bound with Delta RBD (pdb id 8CBF). The heavy chain in orange ribbons, the light chain in red ribbons. (D) The RBD and binding epitope footprint for OMI-42. The binding epitope residues are shown in blue surface. (G) The structure of P5S-1H1 bound with RBD (pdb id 7XS8). The heavy chain in orange ribbons, the light chain in red ribbons. (H) The RBD and binding epitope footprint for P5S-1H1. The binding epitope residues are shown in blue surface. (I) The structure of P5S-2B10 bound with RBD (pdb id 7XSC). The heavy chain in orange ribbons, the light chain in red ribbons. (H) The RBD and binding epitope footprint for P5S-2B10. The binding epitope residues are shown in blue surface.

Although all-atom MD simulations with the explicit inclusion of the glycosylation shield can in principle provide a rigorous assessment of conformational landscape of the SARS-CoV-2 S proteins, such direct simulations are challenging due to the size of a complete SARS-CoV-2 S system embedded onto the membrane and this complexity can often obscure the main molecular determinants of the binding mechanisms and prevents efficient comparative analysis across large number of antibodies. We combined CG-CABS simulations with atomistic reconstruction and additional optimization by adding the glycosylated microenvironment. CG-CABS trajectories were subjected to atomistic reconstruction and refinement. Using dynamic ensembles of the antibody complexes and systematic mutational scanning of the RBD and antibody residues we characterize patterns of mutational sensitivity and compute mutational scanning heatmaps to identify binding hotspot centers and characterize escape mutations. We also employ the Molecular Mechanics/Generalized Born Surface Area (MM-GBSA) approach for rigorous binding affinity computations of the antibody complexes with the S-RBD protein and residue-based energy decomposition. Using these approaches, we examine the dynamic and energetic determinants by which broadly potent class I antibodies can evade immune resistance and dissect energetic mechanisms underlying a uniquely broad neutralization capacity of BD55-1205, which demonstrates extraordinary neutralization breadth against all existing variants as well as prospective variants with mutations within its targeting epitope. We show that despite sharing similar epitopes and interaction footprints, BD55-1205, BD-604, P5S-2B10, and P5S-1H1 induce specific dynamic signatures in the RBD. BD55-1205 stands out for its ability to cause larger stabilization and immobilization of the RBD interface, contributing to its increased stability and stronger binding interactions. In contrast, BD-604 allows for greater flexibility in certain regions, potentially reducing its resilience to mutations. P5S-2B10 and P5S-1H1 exhibit intermediate behavior, balancing stability and flexibility.

We provide a detailed analysis comparing OMI-42, P5S-1H1, P5S-2B10, and BD-604, focusing on their epitope characteristics, binding mechanisms, mutational resilience, and neutralization breadth. The results show that BD55-1205 demonstrates a high degree of ACE2 mimicry, targeting conserved positions which are critical for ACE2 engagement and makes interactions with RBD backbone atoms. The results highlight trade-offs between epitope breadth, binding specificity, and adaptability to mutations. We show that BD55-1205 with broad epitope coverage and distributed hotspot mechanisms, exhibit superior resilience to mutations. In contrast, BD-604 and OMI-42 antibodies featuring more localized binding mechanisms, are more vulnerable to escape mutations at key positions. P5S-1H1 and P5S-2B10 occupy an intermediate position, balancing specificity and adaptability but lacking the robustness and breadth of binding hotspots observed for BD55-1205.

The mutational scanning and MM-GBSA analyses underscore the diversity of binding mechanisms employed by different antibodies, reflecting their unique structural and chemical complementarity with the RBD. The results of this study underscore the fact that similar binding mechanisms for class I antibodies co-exist with subtle interaction differences that could determine the unique neutralization profile. Broad epitope coverage, reliance on multiple binding hotspot residues, and adaptability to mutations are key factors that determine BD55-1205 antibody resilience to immune escape. We show that the evolutionary trade-offs shaping the virus ability to balance immune evasion and ACE2 binding are energetically nuanced and context-dependent. These trade-offs are influenced not only by the specific mutations present the viral genome but also by the characteristics of the antibodies themselves.. This antibody-dependent variability adds another layer of complexity to understanding how SARS-CoV-2 continues to adapt under selective pressures imposed by both natural immunity and vaccination.

## Materials and Methods

### Coarse-Grained Molecular Simulations and Atomistic Reconstruction of Equilibrium Ensembles

Coarse-grained (CG) models are computationally effective approaches for simulations of large systems over long timescales. We employed CABS-flex approach that efficiently combines a high-resolution coarse-grained model and efficient search protocol capable of accurately reproducing all-atom MD simulation trajectories and dynamic profiles of large biomolecules on a long time scale [75–80]. In this high-resolution model, the amino acid residues are represented by Cα, Cβ, the center of mass of side chains and another pseudoatom placed in the center of the Cα-Cα pseudo-bond. In this model, the amino acid residues are represented by Cα, Cβ, the center of mass of side chains and the center of the Cα-Cα pseudo-bond. The CABS-flex approach implemented as a Python 2.7 object-oriented standalone package [75–80] was used in this study to allow for robust conformational sampling proven to accurately recapitulate all-atom MD simulation trajectories of proteins on a long time scale. Conformational sampling in the CABS-flex approach is conducted with the aid of Monte Carlo replica-exchange dynamics and involves local moves of individual amino acids in the protein structure and global moves of small fragments. The default settings were used in which soft native-like restraints are imposed only on pairs of residues fulfilling the following conditions : the distance between their *C*α atoms was smaller than 8 Å, and both residues belong to the same secondary structure elements. The crystal and cryo-EM structures of the Omicron antibody-RBD complexes are obtained from the Protein Data Bank [81] and the following systems were used in this study : the crystal structure of BD55-1205 with XBB.1.5 RBD (pdb id 8XE9), class I antibodies OMI-42 with Delta RBD (pdb id 8CBF), OMI-42 bound with Beta RBD (pdb id 7ZR7), BD-604 bound with RBD (pdb id 7CHF, 7CH4), BD-604 bound with BA.2 RBD (pdb id 8HWT) BD-604 bound with BA.4 RBD (8HWS), P5S-1H1 bound with RBD (pdb id 7XS8), and P5S-2B10 bound with RBD (pdb id 7XSC). A total of 100 independent CG-CABS simulations were performed for each of the systems studied. In each simulation, the total number of cycles was set to 10,000 and the number of cycles between trajectory frames was 100. MODELLER-based reconstruction of simulation trajectories to all-atom representation provided by the CABS-flex package was employed to produce atomistic models of the equilibrium ensembles for studied systems [82]. The CG conformational ensembles of the antibody-RBD complexes were also subjected to all-atom reconstruction using PULCHRA method [83] and CG2AA tool [84]. The all-atom conformations were additionally optimized using the 3Drefine method [85] that utilizes atomic-level energy minimization with composite physics and knowledge-based force fields.

### Binding Free Energy Computations: Mutational Scanning Profiling and Analysis

We conducted mutational scanning analysis of the binding epitope residues for the S RBD-antibody complexes. Each binding epitope residue was systematically mutated using all substitutions and corresponding protein stability and binding free energy changes were computed. BeAtMuSiC approach [86–88] was employed that is based on statistical potentials describing the pairwise inter-residue distances, backbone torsion angles and solvent accessibilities, and considers the effect of the mutation on the strength of the interactions at the interface and on the overall stability of the complex. The binding free energy of protein-protein complex can be expressed as the difference in the folding free energy of the complex and folding free energies of the two protein binding partners:

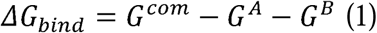

The change of the binding energy due to a mutation was calculated then as the following:

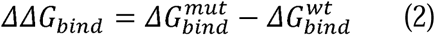

We leveraged rapid calculations based on statistical potentials to compute the ensemble-averaged binding free energy changes using equilibrium samples from simulation trajectories. The binding free energy changes were obtained by averaging the results over 1,000 and 10, 000 equilibrium samples for each of the systems studied.

### Binding Free Energy Computations

We calculated the ensemble-averaged changes in binding free energy using 1,000 equilibrium samples obtained from simulation trajectories for each system under study. Initially, the binding free energies of the RBD-antibody complexes were assessed using the MM-GBSA approach [89,90]. Additionally, we conducted an energy decomposition analysis to evaluate the contribution of each amino acid during the binding of RBD to antibodies [91,92].

The binding free energy for the RBD-Antibody complex was obtained using:

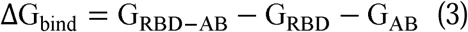

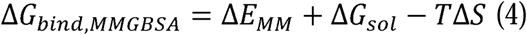

where ΔE_MM_ is total gas phase energy (sum of ΔEinternal, ΔEelectrostatic, and ΔEvdw); ΔGsol is sum of polar (ΔGGB) and non-polar (ΔGSA) contributions to solvation. Here, G _RBD–ANTIBODY_ represent the average over the snapshots of a single trajectory of the complex, G_RBD_ and G_ANTIBODY_ corresponds to the free energy of RBD and antibody respectively. The polar and non-polar contributions to the solvation free energy is calculated using a Generalized Born solvent model and consideration of the solvent accessible surface area [93]. MM-GBSA is employed to predict the binding free energy and decompose the free energy contributions to the binding free energy of a protein–protein complex on per-residue basis. The binding free energy with MM-GBSA was computed by averaging the results of computations over 1,000 samples from the equilibrium ensembles. First, the computational protocol must be selected between the “single-trajectory” (one trajectory of the complex), or “separate-trajectory” (three separate trajectories of the complex, receptor and ligand). To reduce the noise in the calculations, it is common that each term is evaluated on frames from the trajectory of the bound complex. In this study, we choose the “single-trajectory” protocol, because it is less noisy due to the cancellation of intermolecular energy contributions. This protocol applies to cases where significant structural changes upon binding are not expected. Hence, the reorganization energy needed to change the conformational state of the unbound protein and ligand are also not considered. Entropy calculations typically dominate the computational cost of the MM-GBSA estimates. Therefore, it may be calculated only for a subset of the snapshots, or this term can be omitted [94,95]. In this study, the entropy contribution was not included in the calculations of binding free energies of the RBD-antibody complexes because the entropic differences in estimates of relative binding affinities are expected to be small owing to small mutational changes and preservation of the conformational dynamics [94,95]. MM-GBSA energies were evaluated with the MMPBSA.py script in the AmberTools21 package [96] and gmx_MMPBSA, a new tool to perform end-state free energy calculations from CHARMM and GROMACS trajectories [97].

## Results

### Structural Analysis of the RBD Complexes

The structural analysis of binding epitopes for Class I antibodies BD55-1205, BD-604, OMI-42, P5S-1H1, and P5S-2B10 reveals fundamental similarities and yet some subtle differences that could underpin their neutralization breadth, adaptability to mutations, and competition with ACE2 for receptor binding (Figure 2). These antibodies target the apical head of the RBD a highly conserved region critical for ACE2 engagement, ensuring effective blocking of viral entry. However, their epitope coverage, interaction networks, and reliance on specific residues differ thereby shaping their functional profiles and resilience to immune escape. First, we examined and compared the structural organization and composition of epitopes for class I antibodies. Both BD55-1205 and BD-604 exhibit extensive epitope coverage within the RBD, with overlapping regions that include key residues such as Y421, Y453, L455, F456, Y473, A475, G476, N487, Y489, Q493, and H505 (Figure 2A-D, Supporting Information, Tables S2,S3). These residues are distributed across multiple structural elements, including the receptor-binding motif (RBM), hydrophobic core, and receptor-binding ridge (Figure 2A-D, Supporting Information, Tables S2,S3). The epitope of BD55-1205 spans over 20 residues, forming an extensive patch along the receptor-binding ridge, with notable contiguous stretches such as 454– 460, 486–494, 498-505 and (Figure 2A,B, Supporting Information, Table S2). The structural analysis of binding epitopes for Class I antibodies BD55-1205, BD-604, OMI-42, P5S-1H1, and P5S-2B10 reveals fundamental similarities and yet some subtle differences that could underpin their neutralization breadth, adaptability to mutations, and competition with ACE2 for receptor binding (Figure 2).

BD55-1205 engages the largest number of RBD residues, particularly in the RBM region. While the exact set of “key” RBM residues can be nuanced depending on the specific structural study, it is well-established that some of the most consistently highlighted RBM residues involved in ACE2 binding and also frequently mutated in Omicron variants include L455, F456, A475, F486, N487, Y489, F490, L492, Q493, G496, Q498, N501, Y505. All these residues participate in multiple interaction contacts with BD55-1205 (Supporting Information, Table S2). This broad coverage can arguably enhance its resilience to mutations by distributing binding energy across multiple interactions. BD-604 targets a similar but somewhat less extensive epitope (including residues Y421, Y453, L455, F456, Y473, A475, G476, N487, Y489, Q493, and H505) [98,99] but relying more heavily on side-chain contacts rather than backbone-mediated hydrogen bonds, which could limit its adaptability to mutational changes in the RBD (Figure 2C,D, Supporting Information, Table S3). A key distinction between BD55-1205 and BD-604 lies in their reliance on backbone versus side-chain interactions. BD55-1205 forms numerous backbone-mediated hydrogen bonds with RBD residues such as L455, R457, K458, and Q474, enhancing its adaptability to mutations outside the primary epitope (Supporting Information, Table S2). This extensive network of polar interactions, combined with hydrophobic contributions from residues lG416, Y453, L455, and F456, results in a highly stable and resilient interaction. In particular, hydrogen bonds are formed between the backbone atoms of 9 RBD residues (L455, R457, K458, Q474, A475, G476, S490, L492, G502) and HCDR (T28, R31, N32, Y33, P53, R102) [62]. Importantly, BD55-1205 has three unique residues in its HCDRs that make contacts with the RBD backbone atoms: R31 on HCDR1, P53 on HCDR2, and R102 on HCDR3. These specific BD55-1205 positions introduce additional polar interactions with F456, K458, Y473, Q474, A475, G476 [62]. OMI-42 shares a majority of epitope residues with BD55-1205 and BD-604 but demonstrates a slightly narrower but more focused epitope [100,101] (Figure 2E,F, Supporting Information, Table S4). Shared residues between BD55-1205 and OMI-42, such as 453–460 and 473–477, form part of the hydrophobic core and contribute to polar interactions at the receptor-binding ridge (Supporting Information, Tables S2,S4). However, OMI-42 involves only some of the RBM residues in the region 486-494 and unlike BD55-1205, OMI-42 does not make noticeable contacts with F486, F490, L492, S494, G496 (Supporting Information, Table S4).

P5S-2B10 and P5S-1H1 exhibit a nearly identical binding pose to the top face of the RBD, mimicking the binding mode of ACE2 [102] (Figure 2G-J). The binding epitope residues for P5S-1H1 is formed by residues R403, T415, G416, K417, D420, Y421, Y453, L455, F456, R457, K458, N460, Y473, Q474, A475, G476, F486, N487, Y489, Q493, Y495, Q496, Q498, T500, N501, G502, V503 and Y505 (Supporting Information, Table S5). As a result, P5S-1H1 share considerable similarities with BD-604 and BD55-1205 targeting residues within hydrophobic core and mimicking interactions with RBM region including residues 473-477, 484-490, 493-505 (Supporting Information, Table S5). The binding epitope residues for P5S-2B10 consists of residues R403, T415, G416, K417, D420, Y421, Y453, L455, F456, R457, K458, N460, Y473, A475, G476, F486, N487, Y48, Q493, N501, G502 and Y505 (Supporting Information, Table S6). Both P5S-2B10 and P5S-1H1 emphasize residues Y421, Y453, and Y473, which are part of the hydrophobic core and critical for stabilizing the interaction . Central RBM residues, including 415–417 and 420–421, form direct hydrogen bonds and van der Waals contacts, stabilizing the antibody-RBD interface. BD55-1205, BD-604, P5S-2B10, and P5S-1H1 target residues within the RBM and the receptor-binding ridge, which are critical for ACE2 binding. The stretch of residues 453–460 forms a hydrophobic core critical for binding stability of all examined class I antibodies. Among these residues L455 and F456 are particularly important, as they form backbone-mediated hydrogen bonds that enhance adaptability to mutations. Residues 459 and 460 function as hinge points, anchoring the RBD in the complex and contributing to overall stability.

### Coarse-Grained Simulations and Atomistic Reconstruction of the Conformational Ensembles for RBD Complexes with Class I Antibodies

We performed multiple CG-CABS simulations of the SARS-CoV-2 S RBD-antibody complexes followed by all-atom reconstruction of trajectories to examine how structural plasticity of the RBD regions can be modulated by binding of antibodies. The root-mean-square fluctuation (RMSF) profiles provide a detailed view of the dynamic behavior of RBD residues upon antibody binding, highlighting both shared features and notable differences among the antibodies. The primary objective of this study was to investigate the dynamic and energetic contributions of RBD residues, as these residues play a pivotal role in mediating interactions with neutralizing antibodies. However, the flexibility of antibody residue within the complementarity-determining regions (CDRs)—could also influence the overall binding mechanism (Supporting Information, Figure S4). The RMSF profiles revealed generally similar dynamics behavior for RBD residues across all antibodies, reflecting the conserved structural core of the RBD. The central β-sheet and α-helices of the RBD (residues 350–360, 375–380, 394–403) exhibit low RMSF values, indicating minimal flexibility across all complexes (Figure 3). These regions are critical for maintaining the overall structural integrity of the RBD. BD55-1205 induces significant stabilization in RBD regions such as 420–435, 450–475, and 490–505, where residues experience markedly smaller fluctuations compared to other antibodies (Figure 3A). This immobilization locks the RBD into a rigid conformation, reducing ACE2 engagement and enhancing neutralization efficacy (Supporting Information, Figure S4A).

**Figure 3.**
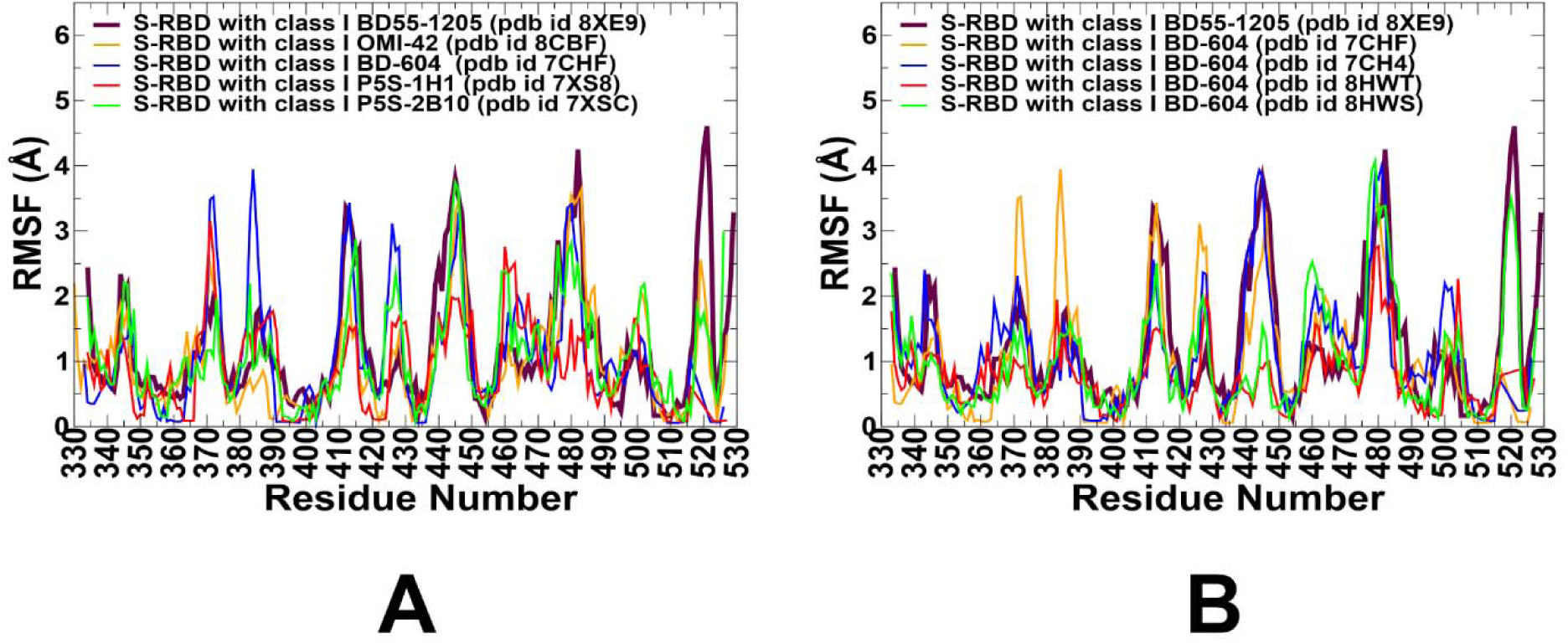
Conformational dynamics profiles obtained from CG-CABS simulations and atomistic reconstruction of the RBD-antibody complexes. (A) The RMSF profiles for the RBD residues obtained from simulations of the S-RBD complexes with class I antibodies : BD55-1205 with XBB.1.5 RBD, pdb id 8XE9 (in maroon thick lines), OMI-42 with Delta RBD, pdb id 8CBF (in orange lines), BD-604 with RBD, pdb id 7CHF/7CH4 (in blue lines), P5S-1H1 with RBD, pdb id 7XS8 (in red lines), and P5S-2B10 with RBD, pdb id 7XSC (in green lines). (B) The RMSF profiles for the RBD residues obtained from simulations of RBD with BD55-1205 (in maroon lines) and different structures of BD-604 antibody with the S-RBD complexes : BD-604 bound with RBD, pdb id 7CHF (in orange lines), BD-604 bound with RBD, pdb id 7CH4 (in blue lines), BD-604 bound with BA.2 RBD, pdb id 8HWT (in red lines), BD-604 bound with BA.4 RBD, pdb id 8HWS (in green lines).

We simulated several structures of BD-604 complexes to obtain a more diverse assessment of the RBD mobility induced by this antibody (Figure 3B, (Supporting Information, Figure S4B). The RMSF profiles suggested that similar to BD55-1205, BD-604 can also cause significant stabilization of the RBD, including regions 375–390. The interfacial RBD positions involved in contacts (residues 490–505) also exhibit reduced flexibility, consistent with their role in stabilizing the antibody-RBD interaction (Figure 3B). The results also revealed important differences, particularly highlighting the subtle specifics of BD55-1205 and BD-604 dynamics. BD-604 binding can enable somewhat larger fluctuations in residues 450-475 suggesting that despite similar binding epitopes and interaction footprints, these class I antibodies can induce specific dynamic signatures. This is consistent with BD-604 exhibiting adaptability to substitutions at specific positions in its complementarity-determining regions (CDRs), such which interact with RBD residues Y473, Q474, A475, G476 [98,99]. Importantly, the results suggest that BD55-1205 binding can cause the larger stabilization and immobilization of the RBD interface which can manifest in the increased RBD stability and stronger binding interactions.

For OMI-42, we found greater stabilization in regions 380-400 and 450-475. This stabilization likely reduces the flexibility of the 470–490 loop, enhancing its ability to block ACE2 binding (Figure 3A, Supporting Information, Figure S4C). P5S-2B10 and P5S-1H1 exhibit intermediate behavior between BD55-1205 and BD-604 as these antibodies do not induce the same level of immobilization as BD55-1205 (Figure 3A, Supporting Information, Figure S4D,E). Regions such as 490–505 show moderate fluctuations, indicating partial stabilization of the RBD interface. Intermediate stabilization suggests that P5S-2B10 and P5S-1H1 maintain a balance between flexibility and rigidity, allowing them to adapt to certain mutations while retaining binding efficacy (Figure 3A). The 470–490 loop of the RBM motif is a highly dynamic region of the RBD, located near the ACE2-binding interface and implicated in stabilizing the RBD-ACE2 interaction. Conformational ensembles obtained from simulations all complexes (Supporting Information, Figure S4) suggested the RBM regions exhibits elevated flexibility, though subtle differences are observed depending on the antibody. BD55-1205 can induce stabilization of 490– 505 indirectly restricts the flexibility of the 470–490 loop, locking the RBD into a rigid conformation. BD-604 may induce more flexibility in 375–390 and 400–420 along with the greater mobility in the 470–490 loop making it potentially less effective at blocking ACE2 interactions (Figure 3A).

To summarize, the results highlight subtle but important distinctions in the dynamic signatures of the RBD induced by class I antibodies. Binding of BD55-1205 stands out owing to significant stabilization of key RBD regions and suggests that BD55-1205 locks the RBD into a rigid conformation, enhancing stability and strengthening binding interactions. BD-604 allows greater fluctuations and induces a less constrained RBD conformation. P5S-2B10 and P5S-1H1 antibodies exhibit intermediate behavior, stabilizing the RBD to some extent but not achieving the same level of immobilization as BD55-1205. This balance between flexibility and rigidity allows these antibodies to adapt to mutations while retaining binding efficacy. The conformational dynamics analysis highlights the nuanced differences in how BD55-1205, BD-604, OMI-42, and P5S-2B10 interact with RBD. BD55-1205 stands out for its significant stabilization, broad epitope coverage, and resilience to mutations, making it highly effective against evolving variants. BD-604, OMI-42, and P5S-2B10 exhibit intermediate behaviors, balancing flexibility and rigidity but lacking the robustness of BD55-1205.

### Mutational Profiling of Antibody-RBD Binding Interactions Interfaces Reveals Molecular Determinants of Immune Sensitivity and Emergence of Convergent Escape Hotspots

Using the conformational ensembles of the RBD-antibody complexes, we embarked on structure-based mutational analysis of the S protein binding with antibodies . To provide a systematic comparison, we constructed mutational heatmaps for the RBD interface residues of the S complexes with class I antibodies. Mutational scanning of BD55-1205 binding revealed a large binding interface and identified multiple distributed hotspot residues that anchor the interaction, including: R403, N405, T415, Y421, Y453, L455, F456, A475, G476, N487, S490, Y501, H505 (Figure 4A,C,D). The large binding interface and multiple hotspot anchors for BD55-1205 distinguishes it from other Class I antibodies. This unique characteristic is reflected in the mutational scanning heatmap showing that strikingly most of the RBD binding interface residues in the complex with BD55-1205 emerged as relevant binding hotspots (Figure 4A).

**Figure 4.**
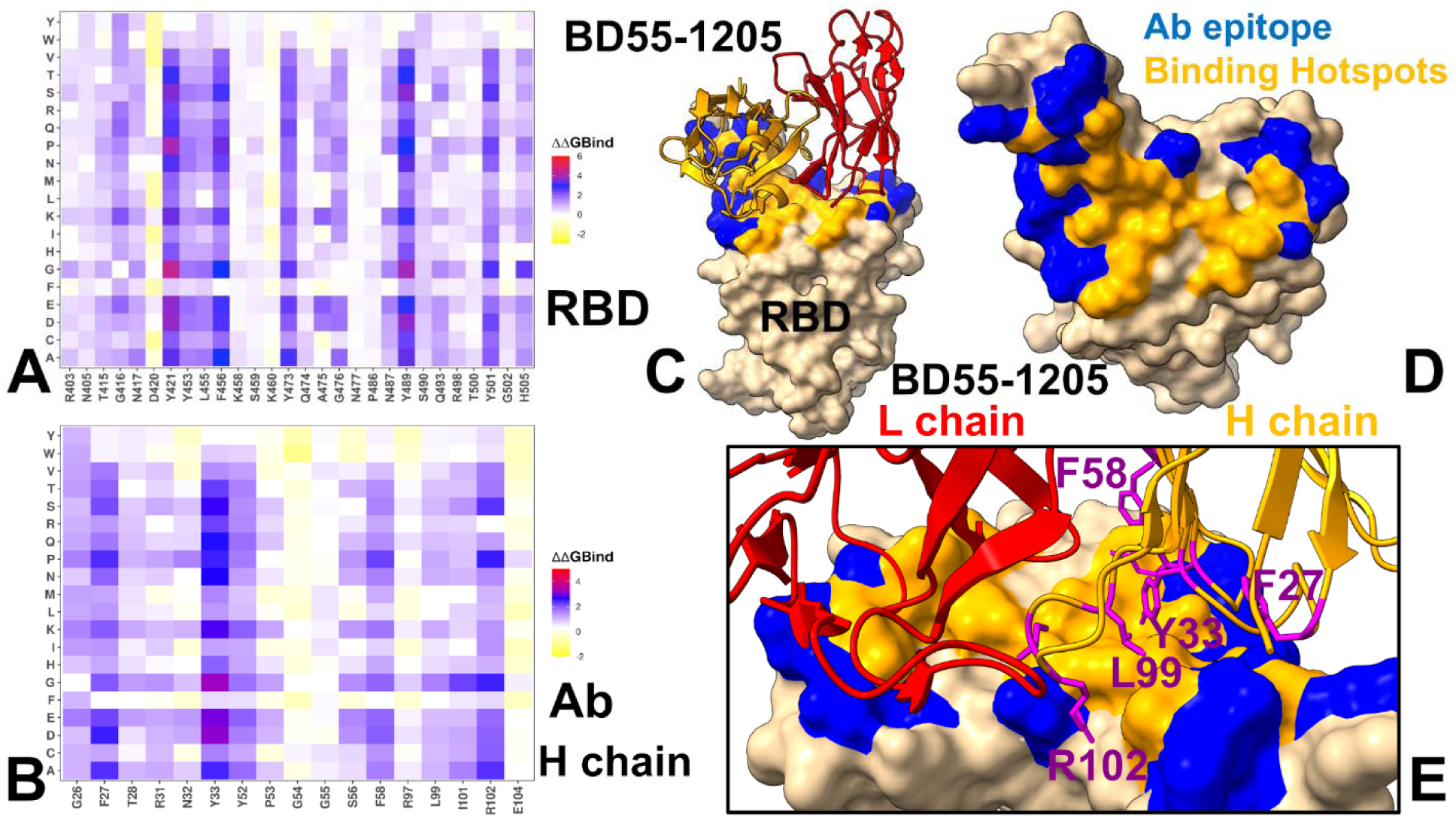
Ensemble-based dynamic mutational profiling of the RBD intermolecular interfaces in the RBD complex with BD55-1205 (A,B). The mutational scanning heatmaps are shown for the interfacial RBD residues (A) and interfacial heavy chain residues of BD55-1205 (B). The heatmaps show the computed binding free energy changes for 20 single mutations of the interfacial positions. The standard errors of the mean for binding free energy changes using randomly selected 1,000 conformational samples (0.11-0.16 kcal/mol) obtained from the atomistic trajectories. (C) The structure of BD55-1205 bound to RBD. The heavy chain of BD55-1205 is in orange ribbons, and light chain is in red ribbons. The binding epitope is shown in blue surface and the positions of the RBD binding energy hotspots R403, N405, T415, Y421, Y453, L455, F456, A475, G476, N487, S490, Y501, H505 are shown in orange-colored surface. (D). RBD from the complex with BD55-1205. The binding epitope residues are in blue surface and the binding interfacial RBD hotspots are in orange surface. (E) A closeup of the binding interface contacts of the BD55-1205 hotspots from the heavy chain F27, Y33, F58, L99 and R102. The heavy chain is in orange ribbons, light chain in red ribbons. The BD55-1205 hotspots F27, Y33, F58, L99 and R102 are shown in magenta sticks and annotated.

Residues T415, Y421, Y453, L455, F456, A475, G476, N487, S490, Y501, H505 form a critical hydrophobic core within the RBM, stabilizing the interaction through van der Waals forces and hydrophobic packing. Y41, Y353, L455, F456 form the central anchoring hotspot cluster and residues Y501, H505 residues function as anchors, providing additional stability to the complex through both polar and hydrophobic interactions. Only several interfacial sites (D420, K458, S459, K460, N477 and P486) in the XBB.1.5 RBD displayed significant tolerance to mutations due to moderate interactions with BD55-1205 (Figure 4A). Conformational dynamics profiles obtained from CG-CABS simulations and atomistic reconstruction of the RBD-antibody complexes. (A) The RMSF profiles for the RBD residues obtained from simulations of the S-RBD complexes with class I antibodies : BD55-1205 with XBB.1.5 RBD, pdb id 8XE9 (in maroon thick lines), OMI-42 with Delta RBD, pdb id 8CBF (in orange lines), BD-604 with RBD, pdb id 7CHF/7CH4 (in blue lines), P5S-1H1 with RBD, pdb id 7XS8 (in red lines), and P5S-2B10 with RBD, pdb id 7XSC (in green lines). (B) The RMSF profiles for the RBD residues obtained from simulations of RBD with BD55-1205 (in maroon lines) and different structures of BD-604 antibody with the S-RBD complexes : BD-604 bound with RBD, pdb id 7CHF (in orange lines), BD-604 bound with RBD, pdb id 7CH4 (in blue lines), BD-604 bound with BA.2 RBD, pdb id 8HWT (in red lines), BD-604 bound with BA.4 RBD, pdb id 8HWS (in green lines)

Mutational scanning identifies residues critical for both RBD stability and high-affinity binding, and destabilizing mutations highlight “hotspots” that are functionally indispensable for the interactions and RBD stability. We noticed that the largest destabilization changes upon BD55-1205 binding are induced in positions Y421 and Y489 that are part of the hydrophobic core and critical for RBD stability and ACE2 binding (Figure 4A). Mutations in these positions can compromise viability of the SARS-CoV-2 S protein and are not observed among variants. Interestingly, our results also showed that mutations in a critical F456 position including F486P (ΔΔG = 2.36 kcal/mol) and F456L (ΔΔG = 1.54 kcal/mol) and F456V (ΔΔG = 1.86 kcal/mol) can affect RBD stability and binding though not as severely as substitutions at positions Y421 and Y489 (Figure 4A). Y421, Y473 and Y489 are critical hotspots for BD55-1205 stability and binding, underscoring their importance in maintaining structural integrity and binding affinity F456 plays a secondary role in stabilizing the complex, with mutations causing moderate destabilization. However, its contribution to hydrophobic packing is crucial for maintaining overall binding affinity. It was suggested that superior neutralizing ability of BD55-1205 against all variants (including the ones with F456L/V) can be attributed to strong interactions with the RBD in this region. We examined the details of mutational scanning data by profiling both RBD and BD55-1205 residues (Figure 4A,C,D). Mutational scanning of the heavy chain BD55-1205 interfacial residues highlighted the two major hotspots Y33 and R102 (Figure 4B). Y33 is the key hotspot for interactions with F456 and R102 makes contacts with L455. We noticed that heavy chain residues Y33, P53, W94, L99, I101 make contacts with L455 on the RBD, while important heavy chain centers R31, N32, Y33, P53, and L99 make contacts with F456 in the complex (Supporting Information, Figure 4B,E). Interestingly, the largest destabilization mutations in the heavy chain are associated with mutations Y33D/E/K/S/N/Q [62]. Interestingly, the experimental structural studies by Cao and colleagues established that BD55-1205 leverages HCDR (T28, R31, N32, Y33, P53, R102) sites to form interactions with RBD backbone of residues L455, R457, K458, Q474, A475, G476, S490, L492, and G502 but not with F456 (Figure 5) [62]. Our analysis suggests that the interactions between Y33 of heavy chain and RBD hotpots Y421, L455 and F456 are among the strongest contacts.

**Figure 5.**
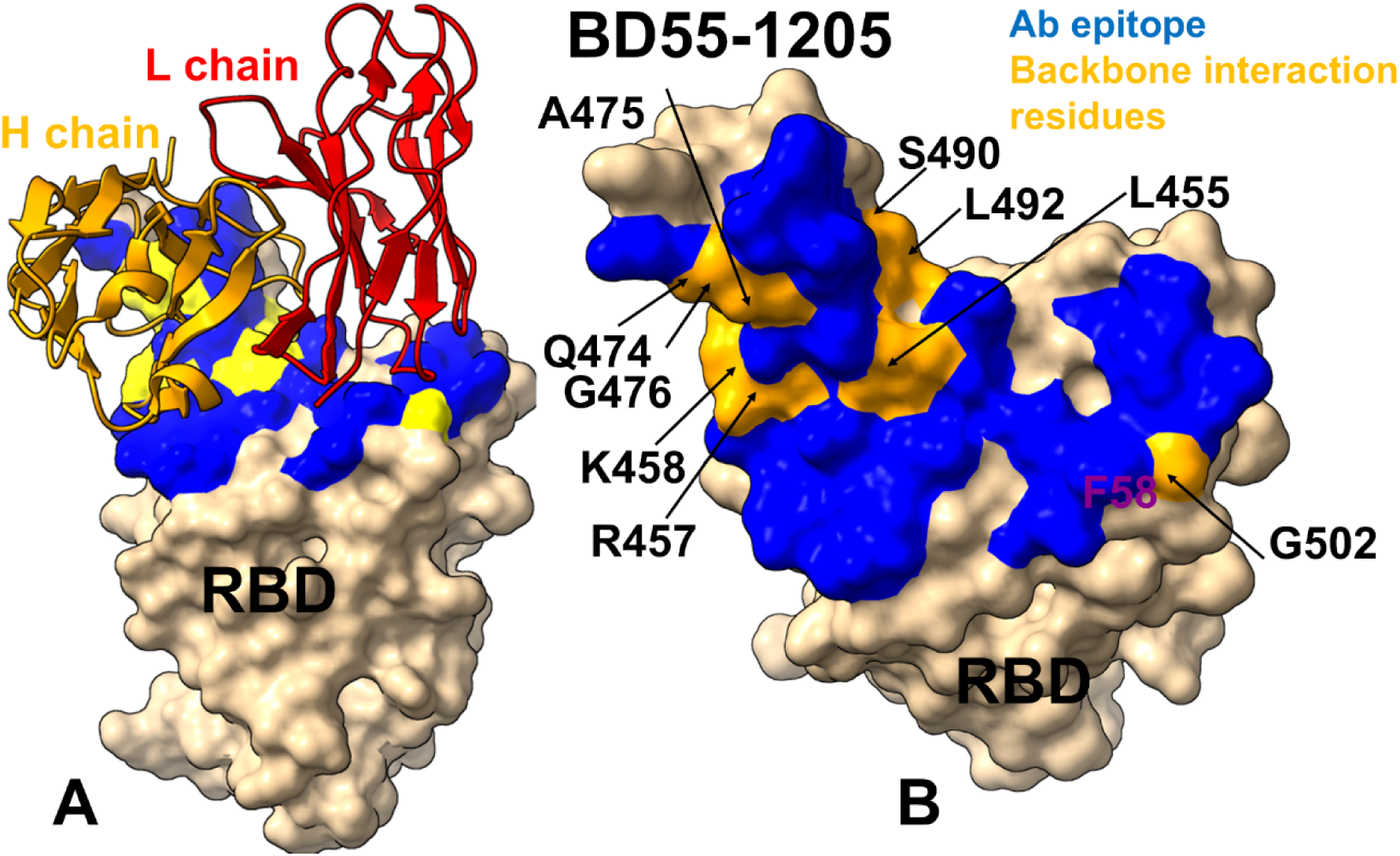
(A)The structure of BD55-1205 bound to RBD. The heavy chain of BD55-1205 is in orange ribbons, and light chain is in red ribbons. The binding epitope is shown in blue surface and the positions of the RBD residues L455, R457, K458, Q474, A475, G476, S490, L492, and G502 that engage their backbone atoms in interfacial interactions with BD55-1205 are shown in yellow surface. (B) RBD from the complex with BD55-1205. The binding epitope residues are in blue surface and the RBD residues L455, R457, K458, Q474, A475, G476, S490, L492, and G502 of backbone interactions with BD55-1205 are shown in orange surface.

The computational predictions are compared against the recent experiments on average antibody escape scores (https://github.com/jbloomlab/SARS2-RBD-escape-calc/tree/main/Cao_data/JN1-evolving-Ab-response/data/DMS/Ab). These experimental data were generated with the escape calculator [103–105] and are reported in the update analysis by Bloom lab (https://jbloomlab.github.io/SARS2-RBD-escape-calc/) that included the latest yeast-display DMS data by Cao and colleagues [46]. By comparing the predicted binding free energy changes obtained from mutational scanning of BD55-1205 residues with the DMS-inferred mutational escape scores (Figure 6), we found an overall excellent overall agreement between the predicted and experimental data.

**Figure 6.**
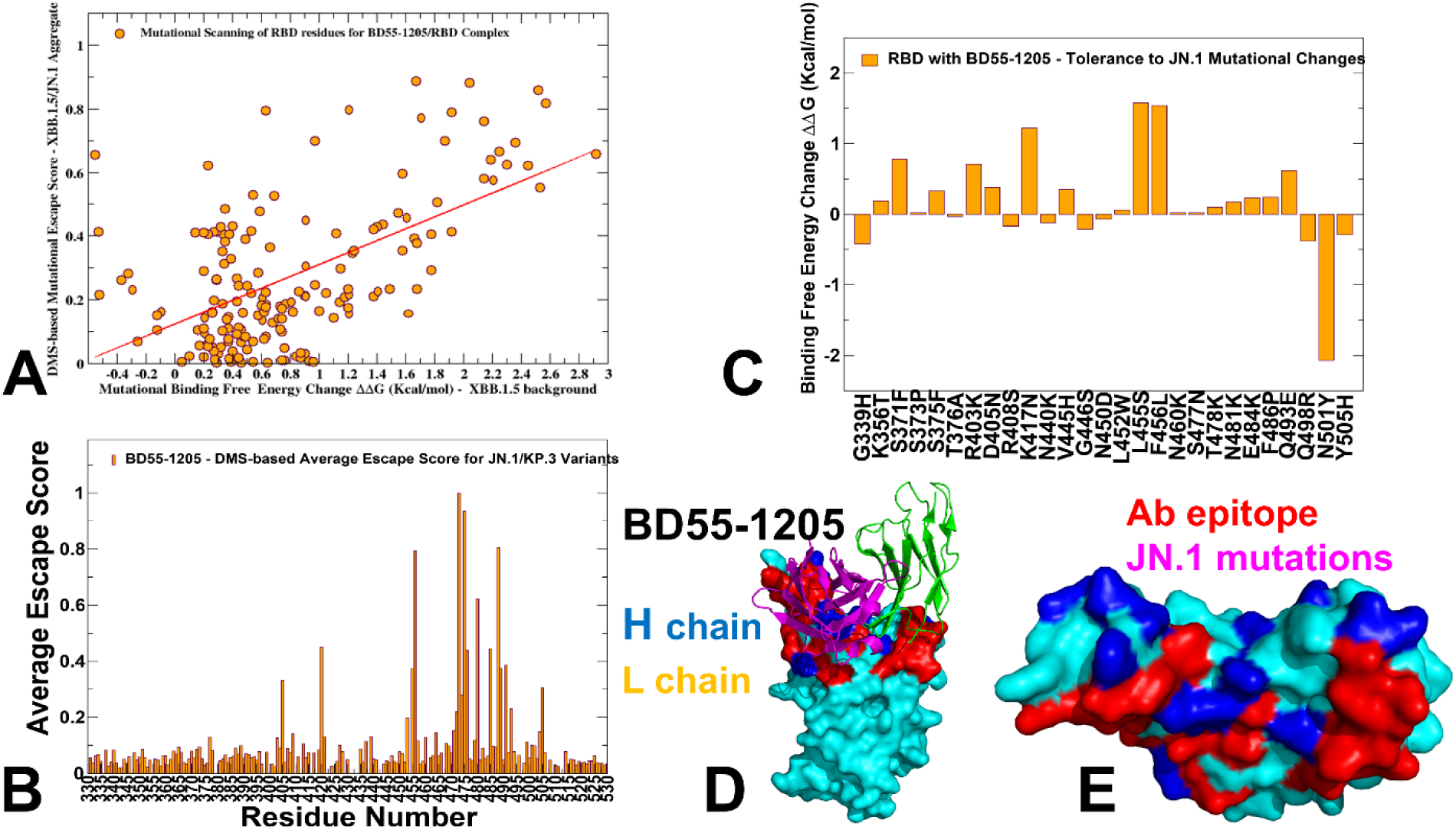
(A) The scatter plot of the predicted binding free energy changes obtained from mutational scanning of BD55-1205 residues versus the experimental DMS-inferred mutational escape scores [62]. (B) The experimental residue-based escape scores of the RBD positions from BD55-1205 antibody. The average escape scores are shown as orange-colored filled bars. (C) Mutational profiling of the S-RBD complex with BD55-1205. The highlighted mutational changes correspond to JN.1 variant mutations. The binding free energy changes are shown in orange-colored filled bars. The positive binding free energy values ΔΔG correspond to destabilizing changes and negative binding free energy changes are associated with stabilizing changes. (D) The structure of BD55-1205 bound to RBD. The heavy chain of BD55-1205 is in blue ribbons, and light chain is in orange ribbons. The binding epitope is shown in red surface and the positions of the JN.1 RBD mutational changes are shown in orange-colored surface. (E). RBD from the complex with BD55-1205. The binding epitope residues are in blue surface and the binding interfacial RBD hotspots are in purple-colored surface.

The Pearson correlation coefficient R=0.57 suggested statistically significant correspondence between the mutational scanning results and the DMS-derived escape scores. The largest binding free energy changes were observed for Y489T, F456G, Y489S, Y473G, F456N, F456P, Y473Q, Y473P and Y473K mutations, while the experimental mutational escape scores displayed the largest escape potential for mutations F456R, A475K, Y473G, F456G, Y473P (Figure 6A). Our predictions accurately predicted important escape mutations in Y489, Y473, A475 and F456 RBD sites. Interestingly, a near-complete loss of neutralization may be indeed associated with F456D, F456E, F456P, F456K, and F456R mutations in various backgrounds but these mutations can also severely reduce ACE2 binding affinity and do not emerge in the evolving variants.

We also examined the experimentally determined average residue-based escape scores that emphasized major escape positions for BD55-1205 and included residues Y473, A475, C488, F456, C480, G485, G476, Y489, and L455 (Figure 6B). While C480 and C488 residues are not involved in the binding interface and are critical for preserving RBD integrity, other critical sites that are located at the binding interface Y473, A475, F456, G476, Y489 and L455 were correctly singled out as mutational hotspots of BD55-1205 binding (Figure 4). In addition, some of the most detrimental mutations revealed by DMS data [46] including Y473A/S/K, A475K/R/H, F456G/N/P are associated with the largest destabilization binding free energies in the computational mutational scanning (Figure 4). Since Y473 and Y489 are fundamentally important for RBD stability and ACE2 binding, the vulnerable sites which may be exploited by virus to escape BD55-1205 are A475 and F456. Both DMS data and our predictions suggested that A475 and F456 mutations may cause localized loss in binding to BD55-1205. In fact, A475V and L455F+F456L mutations emerged in several XBB.1.5 and BA.2.86 lineages enable escape from a broad range of class I antibodies. For instance, LF.7.2.1 demonstrates significantly increased immune evasion compared to XEC, primarily due to the A475V mutation, which enables the virus to escape neutralization by Class 1 antibodies [55]. Hence, the experimentally observed superior neutralizing capacity of BD55-1205 binding [62] can be attributed not to the lack of sensitivity to mutations in A475 and F456 but likely due to a broad and efficiently distributed network of hotspots and interactions that may enable to sustain and partly mitigate the effect of mutations in these positions. The suggested distributed anchoring binding mechanism could contribute to BD55-1205 resilience to mutations. In this mechanism, even if one or two hotspot residues are mutated (L455 and/or F446), the overall binding affinity would not be severely compromised due to the redundancy in stabilizing interactions.

Mutational scanning for BD-604 binding revealed a reduced breath in the distribution of main hotspots, pointing to Y421, Y453, L455, and F456, Y489, R493, Y501, G502 and H505 as main energetic centers which are also critical for ACE2 engagement (Figure 7A). Interestingly, the major destabilization mutations in BD-604 are generally similar to those in BD55-1205 and are associated with the protein stability effects but the extent of destabilization is not as significant as for BD55-1205 (Figure 7A,C,D). We also similarly evaluated mutational profiles for BD-604 and found that the most destabilizing mutations are associated with Y33 and Y102 and G26 (Figure 7B). BD-604 exhibits greater tolerance for mutations of I28, S30, and S31 in HCDR1 that are more adaptable to substitutions. These residues interact with RBD residues Y473, Q474, A475, G476, Y501 and H505 suggesting a broader adaptability to certain mutations (Figure 7E) but weaker cumulative interactions with the RBD residues. These differences may underscore BD55-1205 enhanced binding affinity and mutational resilience compared to BD-604.

**Figure 7.**
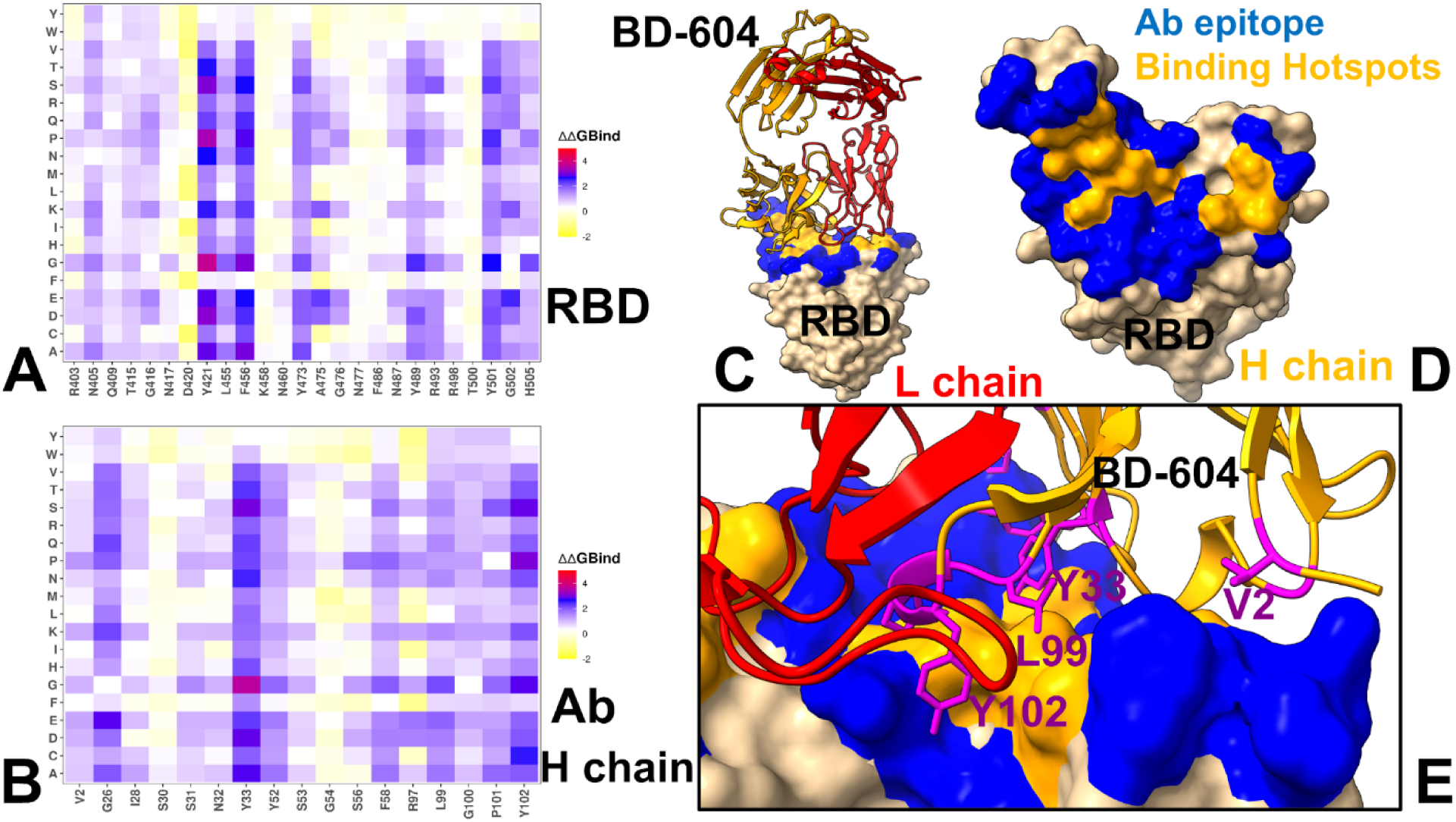
Ensemble-based dynamic mutational profiling of the RBD intermolecular interfaces in the RBD complex with BD-604 (A,B). The mutational scanning heatmaps are shown for the interfacial RBD residues (A) and interfacial heavy chain residues of BD-604 (B). The structure of BD-604 bound with BA.2 RBD (pdb id 8HWT) is used in this analysis. The heatmaps show the computed binding free energy changes for 20 single mutations of the interfacial positions. The standard errors of the mean for binding free energy changes using randomly selected 1,000 conformational samples (0.08-0.14 kcal/mol) obtained from the atomistic trajectories. (C) The structure of BD-604 bound to RBD. The heavy chain of BD-604 is in orange ribbons, and light chain is in red ribbons. The binding epitope is shown in blue surface and the positions of the RBD binding energy hotspots Y421, Y453, L455, and F456, Y489, R493, Y501, G502 and H505 are shown in orange-colored surface. (D). RBD from the complex with BD-604. The binding epitope residues are in blue surface and the binding interfacial RBD hotspots are in orange surface. (E) A closeup of the binding interface contacts of the BD-604 hotspots from the heavy chain V2, Y33, L99 and Y102. The heavy chain is in orange ribbons, light chain in red ribbons. The BD-604 hotspots V2, Y33, L99 and Y102 are shown in magenta sticks and annotated.

Mutational scanning map suggested that RBD positions N417, N460 and F486 are moderately tolerant to substitutions while mutations in L455 and F456 are highly destabilizing (Figure 7A). Due to less extensive interaction network formed by BD-604 could make it vulnerable to mutations in variants BA.4/BA5 and XBB.1.5. Indeed, the neutralizing ability of BD-604 showed considerable reduction in neutralization against BA.4/5, BQ.1.1, and XBB variants, highlighting the fact that mutations K417N, N460K in BA4/BA.5 and V445P, G446S, N460K, F486S, and F490S can reduce BD-604 binding, and these cumulative losses are far more detrimental for BD-604 activity than for BD55-1205. For consistency and comparison, we also built mutational scanning heatmaps for other structures of antibody BD-604 bound with RBD (pdb id 7CHF, 7CH4), BD-604 bound with BA.2 RBD (pdb id 8HWT) BD-604 bound with BA.4 RBD (8HWS), RBD, BA.1 RBD and BA.2 RBD (Supporting Information, Figures S5,S6). The results are very consistent among these complexes, emphasizing critical triad of RBD hotspot positions Y453, L455, F456 as well as RBD residues Y473, Y489, Y501, G502 and H505. To provide a systematic comparison, we also constructed mutational heatmaps for the RBD interface residues of other S-RBD complexes with class I OMI-42 (Supporting Information, Figure S7). OMI-42 is susceptible to A475V and L455F+F456L mutations emergent in several XBB.1.5 and BA.2.86 lineages [106]. The mutational heatmap analysis showed that positions Y421, Y453, L455 and F456 emerged as key escape hotspots with Omi-42 (Supporting Information, Figure S7A,C,D). L455 and F456 positions are located at the epitope of RBD Class 1 antibodies and neutralization assays demonstrated that L455S mutation enables JN.1 to evade Class 1 antibodies. Mutational heatmaps data showed that effectively all modifications in L455, including L455S, can cause considerable loss in antibody binding. A secondary group of escape hotspots for OMI-42 included F486, N487, Y489, and Q493 positions (Supporting Information, Figure S7A,C,D). Mutational profiling of heavy chain residues of OMI-42 showed significant role of Y32, W53, F101, Y109 and Y110 (Supporting Information, Figure S7B,E). Compared to BD55-1205 and another ultrapotent antibody VIR-7229, OMI-42 forms fewer hydrogen bonds in the region near L455/F456 which may explain the markedly reduced neutralizing activity for XBB-descendant and JN.1-descendant variants harboring F456L [63]. Omi-42 binding is abrogated by several substitutions at F456 position in multiple backgrounds [63]. This contrasts with. BD55-1205 which has an exceedingly high barrier to viral resistance as the relative tolerance for epitope diversification is promoted by the extensive contacts with the RBD backbone, P5S-2B10 and P5S-1H1 exhibit a nearly identical binding pose to the top face of the RBD, mimicking the binding mode of ACE2. These antibodies compete directly with ACE2 for binding. Mutational scanning of P5S-1H1 revealed key hotspots Y421, Y453, L455, F456 and H505 while positions K417, Y473, Y489 and N501 may be less sensitive to mutations, reflecting greater conformational mobility of the RBD in this complex (Figure 8A-D). The analysis of P5S-1H1 pointed to Y33, Y52, F58, L99, and Y102 as important hotspot centers (Figure 8B,E).

**Figure 8.**
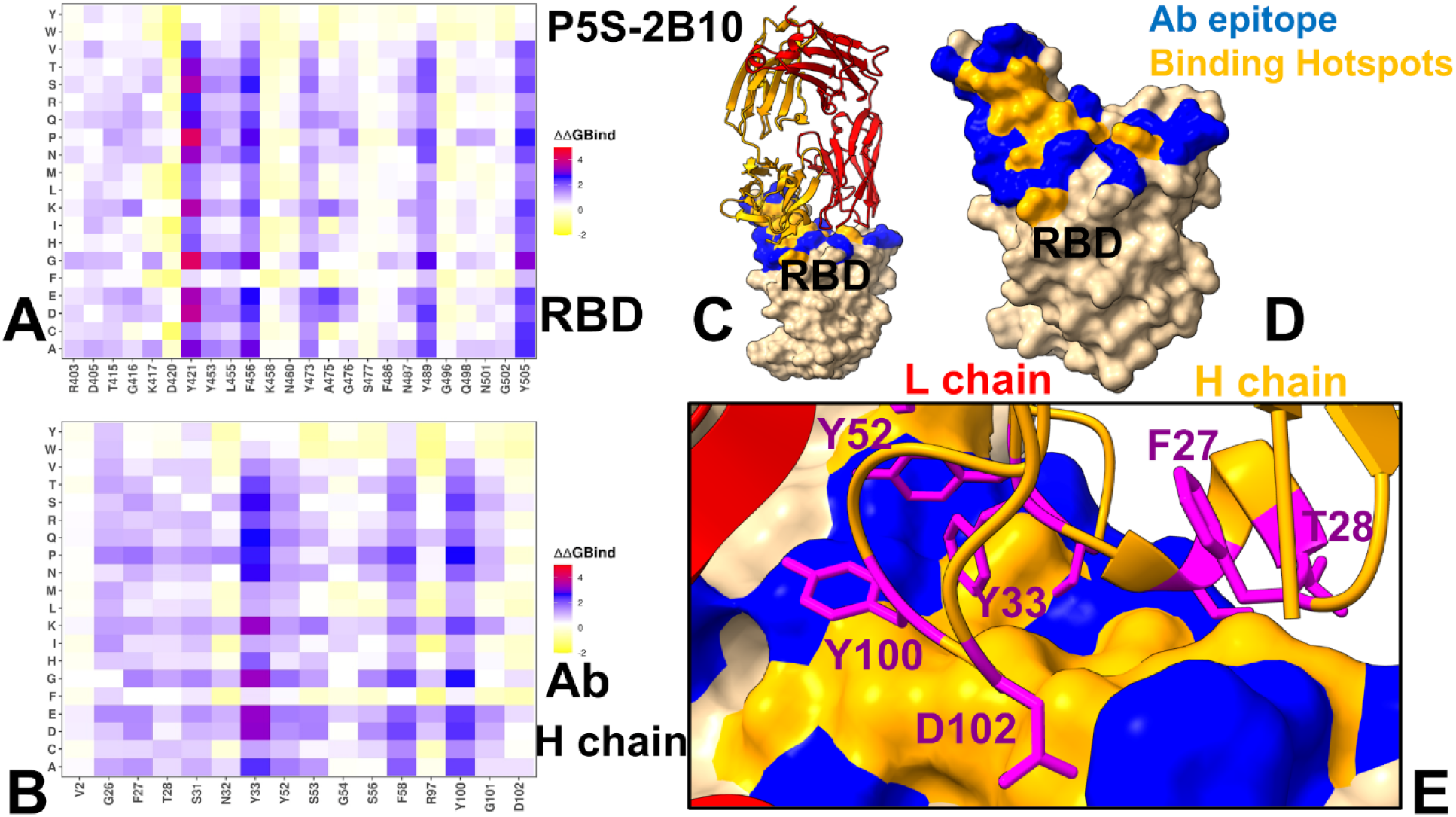
Ensemble-based dynamic mutational profiling of the RBD intermolecular interfaces in the RBD complex with P5S-1H1 (A,B). The mutational scanning heatmaps are shown for the interfacial RBD residues (A) and interfacial heavy chain residues of BD-604 (B). The structure of P5S-1H1 bound with RBD (pdb id 7XS8) is used in this analysis. The heatmaps show the computed binding free energy changes for 20 single mutations of the interfacial positions. The standard errors of the mean for binding free energy changes using randomly selected 1,000 conformational samples (0.17-0.22 kcal/mol) obtained from atomistic trajectories. (C) The structure of P5S-1H1 bound to RBD. The heavy chain of BD-604 is in orange ribbons, and light chain is in red ribbons. The binding epitope is shown in blue surface and the positions of the RBD binding energy hotspots Y421, Y453, L455, F456 and H505 are shown in orange-colored surface. (D). RBD from the complex with P5S-1H1. The binding epitope residues are in blue surface and the binding interfacial RBD hotspots are in orange surface. (E) The binding interface contacts of the P5S-1H1 hotspots from the heavy chain Y33, S53, F58, L99 and Y102. The heavy chain is in orange ribbons, light chain in red ribbons. The P5S-1H1 Y33, S53, F58, L99 and Y102 are in magenta sticks and annotated.

The mutational heatmaps for P5S-2B10 are generally similar and identified Y421, Y453, L455, F456 and H505 as major hotspots (Figure 9A) while the corresponding energetic anchors on the antibody correspond to Y33, F58, Y100 but Y102 position is more tolerant and less energetically important (Figure 9B,E). Strikingly, in all RBD complexes with class I antibodies mutational heatmaps clearly showed that a pair of adjacent residues L455 and F456 corresponding to convergent evolutionary hotspots are also dominant escape hotpots of antibody neutralization. To summarize, despite targeting very similar epitopes and sharing comparable binding modes, BD55-1205, BD-604, OMI-42, P5S-1H1, and P5S-2B10 exhibit subtle yet critical differences in their interaction patterns with the RBD. These distinctions arise from variations in epitope composition, reliance on specific residues, adaptability to mutations, and structural dynamics, collectively shaping their efficacy, breadth, and resilience to viral evolution. The comparison between BD55-1205 and other neutralizing class I antibodies further underscored the most striking difference – a large and broadly distributed footprint of strong hotspots for BD55-1205 in contrast to more localized distribution of major hotspots concentrated in Y421, Y453, L455 and F456 positions.

**Figure 9.**
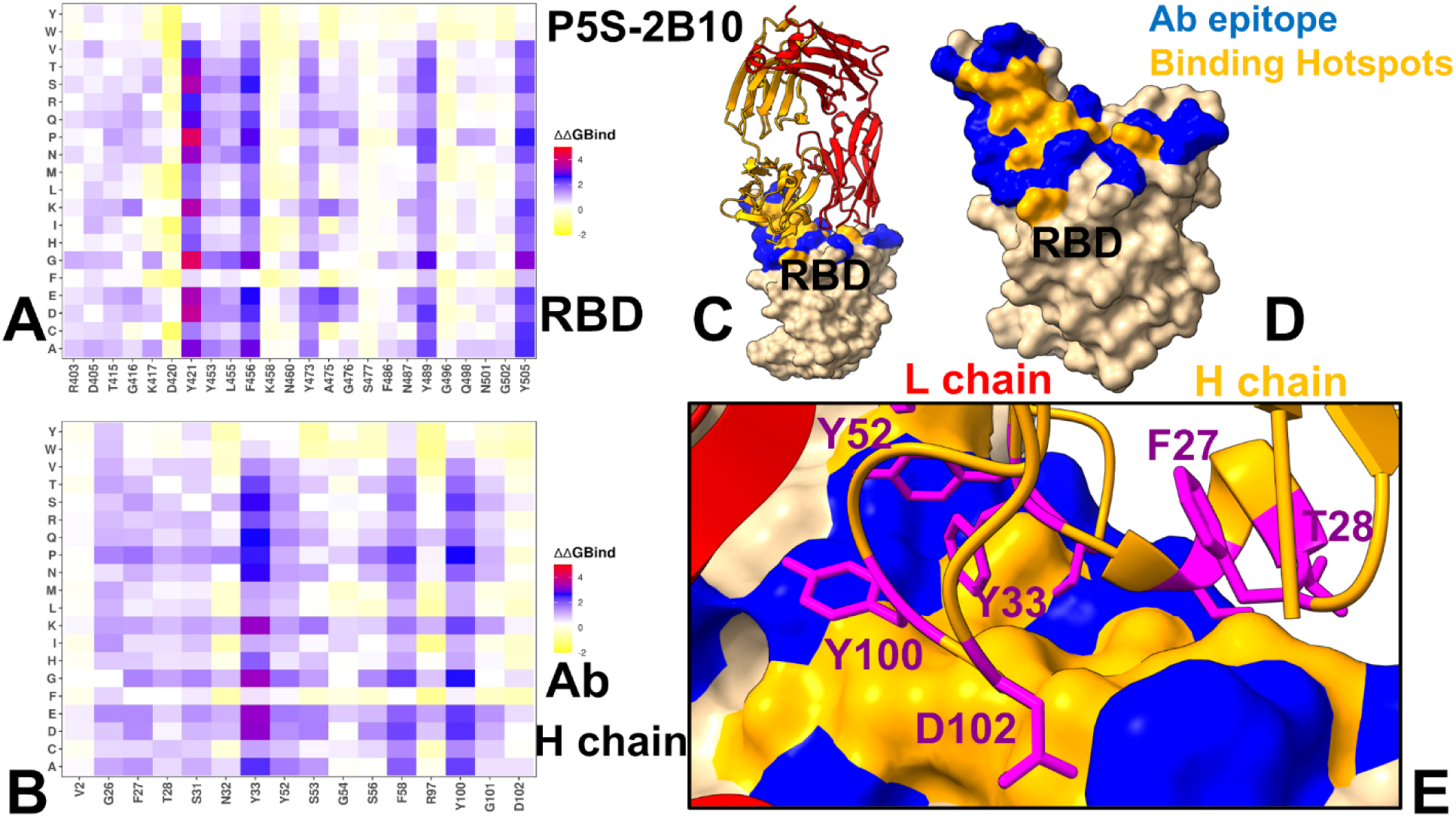
Ensemble-based dynamic mutational profiling of the RBD intermolecular interfaces in the RBD complex with P5S-2B10 (A,B). The mutational scanning heatmaps are shown for the interfacial RBD residues (A) and interfacial heavy chain residues of BD-604 (B). The structure of P5S-2B10 bound with RBD (pdb id 7XSC) is used in this analysis. The heatmaps show the computed binding free energy changes for 20 single mutations of the interfacial positions. The standard errors of the means for binding free energy change using randomly selected 1,000 conformational samples (0.07-0.12 kcal/mol) obtained from atomistic trajectories. (C) The structure of P5S-2B10 bound to RBD. The heavy chain of P5S-2B10 is in orange ribbons, and light chain is in red ribbons. The binding epitope is shown in blue surface and the positions of the RBD binding energy hotspots Y421, Y453, L455, F456 and H505 are shown in orange-colored surface. (D). RBD from the complex with P5S-2B10. The binding epitope residues are in blue surface and the binding interfacial RBD hotspots are in orange surface. (E) A closeup of the binding interface contacts of the P5S-2B10 hotspots from the heavy chain Y33, F27, T28, Y52, Y100, D102. The heavy chain is in orange ribbons, light chain in red ribbons. The P5S-2B10 hotspots Y33, F27, T28, Y52, Y100, D102are shown in magenta sticks and annotated.

The mutational scanning analysis underscores the subtle differences of binding mechanisms employed by class I antibodies targeting remarkably similar epitopes but exerting distinct energetic patterns with the RBD. The results suggested that major drivers of immune escape for this class I antibodies correspond to L455 and F456 sites that undergo mutations in the JN.1, KP.2 and KP.3 variants, enabling evolution through the enhanced immune escape. L455 and F456 consistently emerge as dominant escape hotspots across all antibodies, enabling viral evolution through enhanced immune evasion. BD55-1205 leverages a broad network of backbone-mediated hydrogen bonds with RBD residues and features a more extensive set of strong binding hotspots thereby rescuing potential losses and enhancing adaptability to mutations in commonly shared positions L455 and F456. BD-604, OMI-42, P5S-1H1 and P5S-2B10 antibodies target similar epitope residues and are also similar in the composition of major binding hotspots that are concentrated on L455 and F456 positions making them more vulnerable to specific mutations. The reduced breadth of highly favorable interactions with the RBD as compared to BD55-1205 increases vulnerability of other class I antibodies to cumulative losses caused by multiple mutations, highlighting mechanisms underlying more limited adaptability to evolving variants.

### Binding Energetics and Residue-Based Decomposition Analysis for Class I Antibody-RBD Complexes: Broadly Distributed Footprint of Multiple Binding Hotspots Determines Unique Neutralization Profile of BD55-1205

We utilized conformational equilibrium ensembles derived from CG-CABS simulations to compute the binding free energies for the RBD-antibody complexes using the MM-GBSA method. This approach allowed us to identify key binding hotspots and quantify the roles and synergies of van der Waals and electrostatic interactions in the binding mechanism. Through MM-GBSA analysis we identify the binding energy centers on the RBD and antibodies and examine the binding mechanisms and energetic determinants of immune escape. We analyzed the MM-GBSA results for BD55-1205 binding to the RBD and examined the energetic contributions of key residues and their roles in stabilizing the antibody-RBD complex. The key energetic centers on the RBD are broadly spread and correspond to residues Y489, H505, A475, Y421, L492, Y501, F456, L455, G476 and N487 displaying the largest total binding energies (Figure 10A) The interesting comparative insights are obtained from the analysis of van der Waals interactions where the largest contributions correspond to residues H505, L455, F456, A475, Y421, Y489, N487, and Y473 (Figure 10A). For example, F456 and L455 contribute significantly to hydrophobic interactions, as evidenced by their favorable van der Waals energies (−3.88 kcal/mol and -2.04 kcal/mol, respectively) (Figure 10B). These results are consistent with the mutational scanning analysis of BD55-1205 interactions with the RBD which revealed positions Y421, Y453, L455, F456, A475, G476, N487, S490, Y501, H505 as dominant energetic centers. MM-GBSA results emphasized the role of H505, Y501, F456, A475, and Y489 sites due to their favorable van der Waals forces and hydrophobic packing (Figure 10B). The strongest electrostatic interactions are formed with R403, E471, D467 and N487 residues where R403 and N487 positions contribute favorably through some synergy of the van der Waals and electrostatic contributions (Figure 10C). By exploiting consensus between MM-GBSA and mutational scanning predictions, we found that residues Y473, A475, F456, Y489 and H505 contributing strongly to both van der Waals and electrostatic interactions. Overall, the hydrophobic contacts provide dominant contribution to the total binding energy and particularly favorable and synergistic with other contributions for Y421, F456, Y473, A475, Y489 and H505 positions (Figure 10A-C). Strikingly, our predictions are in excellent agreement with DMS escape profiles (based on XBB.1.5 RBD) for BD55-1205 showing that major immune escape sites are Y473, A475, F456, Y489 [62]. Critical residues Y421 and Y489 are indispensable for RBD stability and ACE2 binding, making them less prone to mutational escape despite their importance in antibody binding. Notably, the DMS data also pointed to N487 and H505 that displayed highly favorable interactions with BD55-1205 (Figure 10A-C). While certain residues like F456 and L455 can be moderately sensitive to mutations, the overall network of interactions provides a higher barrier to resistance compared to antibodies with narrower epitopes. It is important to note that BD55-1205 leverages interactions with the RBD backbone in positions L455, R457, K458, Q474, A475, G476, S490, L492, and G502 [62]. Some of these residues A475, G476 and L492 are among strong energetic centers but the spectrum of binding energy hotspots appeared to be much broader and included also Y421, Y453, F456, Y501, H505 residues. This analysis suggested that the “backbone-interacting” RBD positions could provide a second “line of defense” against BD55-1205 resistance as mutations in convergent evolution sites L455 and F456 may be potentially rescued and compensated by sustainable backbone-based interactions (Figure 10).

**Figure 10.**
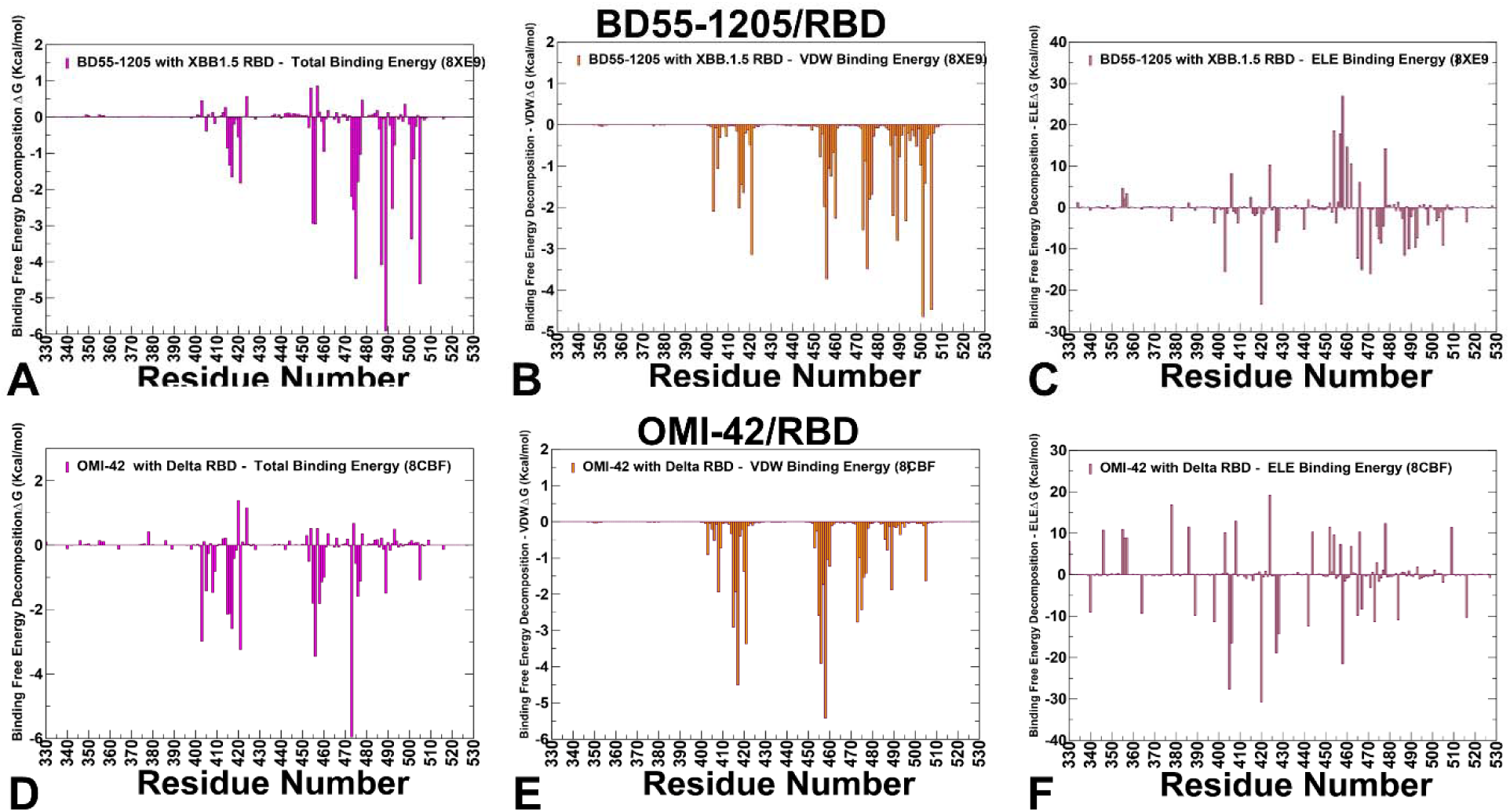
The residue-based decomposition of the binding MM-GBSA energies for BD55-1205 and OMI-42 binding. The residue-based decomposition for BD55-1205/RBD binding shows the total binding energy (A), van der Walls contribution to the total MM-GBSA binding energy (B) and the electrostatic contribution to the total binding free energy (C). MM-GBSA contributions are evaluated using 1,000 samples from the simulations of BD55-1205 complex with RBD, pdb id 8XE9. The residue-based decomposition for OMI-42/RBD binding shows the total binding energy (D), van der Walls contribution to the total MM-GBSA binding energy (E) and the electrostatic contribution to the total binding free energy (F). MM-GBSA contributions are evaluated using 1,000 samples from simulations of OMI-42 complex, pdb id 8CBF. Residue-based binding free energy values are shown for total energies as magenta-colored filled bars, van der Waals contributions as orange-colored bars and electrostatic contributions as light-brown colored bars.

Based on the MM-GBSA analysis and mutational scanning data, we suggest that the unique binding mechanism of BD55-1205 is determined by its broadly distributed network of binding hotspots that may define a high barrier to viral resistance. While some hotspots are important for ACE2 binding and therefore functionally and evolutionary constrained, high tolerance for mutations in L455/F456 region can be enabled through an extensive interaction network of presence of diverse additional RBD hotspots. The MM-GBSA calculations for another class I antibody OMI-42 identify several residues as critical hotspots for OMI-42 binding: Y473, F456, Y421, R403, K417, T415 and G416 where Y473 and F456 dominate the total binding energy (Figure 10D). The breakdown of the binding free energy components revealed that F456, K417, Y421, T415, Y473, L455 and A475 contribute strongly to van der Waals stabilization (Figure 10E), while the electrostatic interactions favor D420, D405, K458, D427 but their total contributions are relatively moderate due to solvation effects (Figure 10F). Overall, the MM-GBSA analysis revealed strong dominant contributions of F456, Y421, T415, Y473, L455 and A475 to binding and may be susceptible to mutations in this positions. Although the binding epitope for OMI-42 is generally similar to BD55-1205, the footprint of major binding hotspots is somewhat different and more localized compared to BD55-1205, making it more susceptible to mutations at critical positions L455, F456, A475 (Figure 10D-F). The results confirmed the experimental finding that KP.2 and KP.3 harboring the F456L mutation can significantly impair the neutralizing activity of Omi-42 [45]. Indeed, the functional experiments showed that Omi-42 is susceptible to A475V and L455F+F456L mutations emerged in several XBB.1.5 and BA.2.86 lineages and may be partly affected by D405N and R408 [100].The recently circulating evasive variants, including HK.3.1, JD.1.1, and JN.1 could efficiently escape OMI-42 due to their escape mutations (L455F + F456L) of HK.3.1 and the additional A475 V mutation carried by JD.1.1 [107,108]. In contrast, BD55-1205 demonstrates superior mutational resilience due to its broad epitope coverage and cumulative binding strength. In summary, the L455 and F456 positions appear to be particularly critical vulnerabilities for OMI-42, with mutations at these sites providing significant escape potential for the virus. The A475V mutation also seems to be an important escape route specifically for this antibody. These differences highlight the trade-offs between epitope breadth, residue dependence, and adaptability to mutations.

MM-GBSA for BD-604 binding emphasized the role of N487, Y505, Y489, A475, G502, Q498, K417, Y421 and N460 residues to the total binding energy (Figure 11A). In particular, the van der Waals interactions were strongest for K417, Y505, Y489, Y421 and L455 (Figure 11B). The electrostatic contacts were favorable K417, R403 and D420 but were often largely offset by unfavorable solvation (Figure 11C). Overall, BD-604 binding is also dependent significant number of hotspots, but their total binding free energies are weaker than the binding energetics of BD55-1205. Additionally, MM-GBSA analysis showed there may be less synergy between hydrophobic and electrostatic contributions in driving binding preferences of major hotspots for BD-604. (Figure 11B,C). For comparison, we also examined MM-GBSA energies using ensemble obtained from simulations of another BD-604 complex with RBD (Figure 11D-F). As may be expected, the results revealed similar patterns, highlighting N487, L455, F456, A475, H505 and Y489 as key binding hotspots (Figure 11D-F). In contrast to BD55-1205, BD-604 exhibits a more localized binding mechanism, focusing its energy on energetically dominant residues such as Y421, Y453, L455, F456, Y489, G502, and H505 that include functionally and evolutionary constrained positions indispensable for RBD stability and ACE2 binding (Y421, Y453, Y489, G502, H505) along with convergent evolution centers L455 and F456. As a result, mutations in L455 and F456 positions could have a more profound effect on BD-604 binding since these modifications can not be fully rescued through “backbone-based” robust interactions and presence of additional hotspot centers. BD-604 also relies more heavily on side-chain contacts rather than backbone-mediated hydrogen bonds, making it less adaptable to structural changes in the RBD. The experimentally measured escape profiles and neutralizing ability of BD-604 showed considerable reduction in neutralization against BA.4/5, BQ.1.1, and XBB variants, highlighting the fact that mutations K417N, N460K in BA4/BA.5 and V445P, G446S, N460K, F486S, and F490S can reduce BD-604 binding [100]. The MM-GBSA breakdown showed that BD-604 binding largely relies on van der Waals forces but exhibited the overall weaker cumulative interactions formed by BD-604 as compared to BD55-1205, thus rendering it less effective against broader antigenic variation.

**Figure 11.**
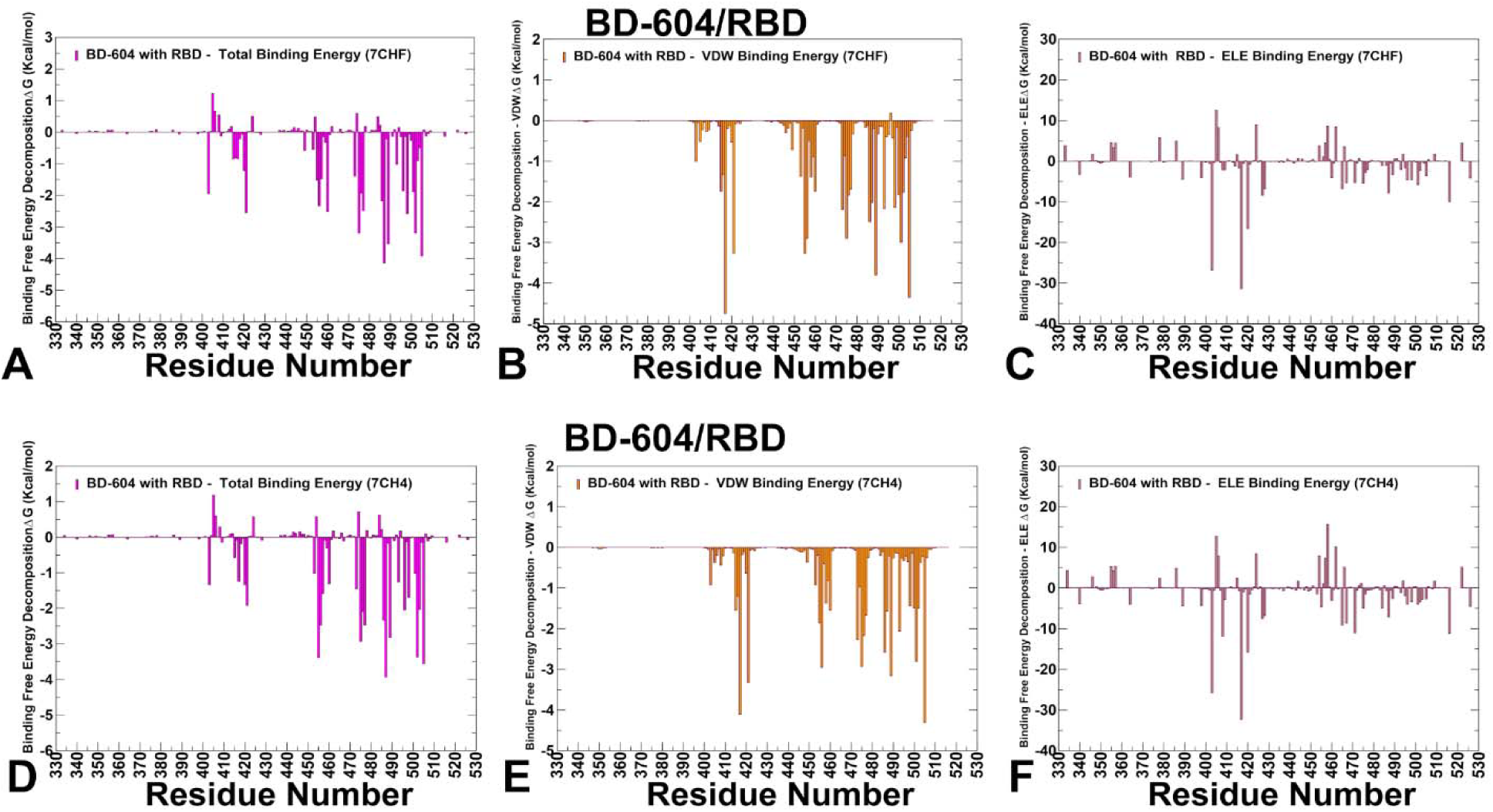
The residue-based decomposition of the binding MM-GBSA energies for BD-604 binding. The residue-based decomposition for BD-604/RBD binding shows the total binding energy (A), van der Walls contribution to the total MM-GBSA binding energy (B) and the electrostatic contribution to the total binding free energy (C). MM-GBSA contributions are evaluated using 1,000 samples from the simulations of BD-604 complex with RBD, pdb id 7CHF. The residue-based decomposition for BD-604/RBD binding shows the total binding energy (D), van der Walls contribution to the total MM-GBSA binding energy (E) and the electrostatic contribution to the total binding free energy (F). MM-GBSA contributions are evaluated using 1,000 samples from simulations of OMI-42 complex, pdb id 7CH4. Residue-based binding free energy values are shown for total energies as magenta-colored filled bars, van der Waals contributions as orange-colored bars and electrostatic contributions as light-brown colored bars.

P5S-1H1 and P5S-2B10 antibodies belong to Class I neutralizing antibodies, which target epitopes overlapping with the ACE2-binding site. MM-GBSA analysis of P5S-1H1 revealed a number of key hotspots including K417, Y505, A475, N487, L455, F486, N501, G502 (Figure 12A). The strongest residues displaying favorable van der Waals contacts are Y505, A475, Y489, Y421, F456 and F486 (Figure 12B). K417 exhibits strong van der Waals and electrostatic interactions while A475, L455 and F486 contribute primarily through van der Waals interactions, stabilizing the hydrophobic core within the receptor-binding motif (RBM) (Figure 12B). Interestingly, unlike BD1205 and BD-604, this antibody binding is less dependent on F456 site but strongly relies on electrostatic contacts with K417 (Figure 12C) and hydrophobic interactions with A475, Y489, Y505 and F486. As a result, mutations in K417, Y505 and F486 seen in various Omicron variants can reduce the neutralizing activity of P5S-1H1 [62]. These results suggested that P5S-1H1 can rely significantly on K417 and could be affected by mutations in K417. Additionally, P5S-1H1 makes important contacts with L455, F486 residues that are mutated in various Omicron variants including XBB.1.5, SLip, FLiRT, and KP.2 variants.

**Figure 12.**
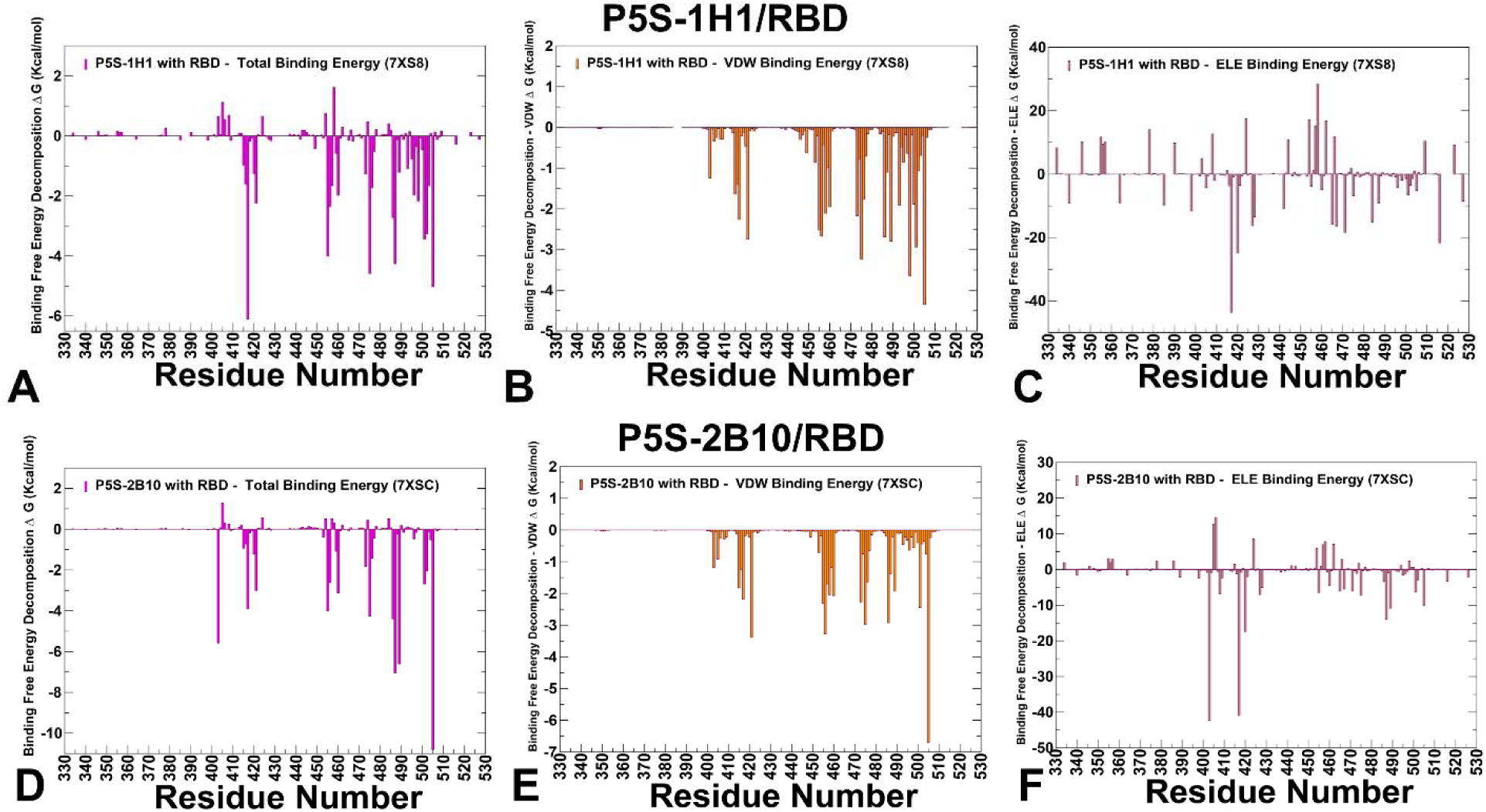
The residue-based decomposition of the binding MM-GBSA energies for P5S-1H1 and P5S-2B10 binding. The residue-based decomposition for P5S-1H1/RBD binding shows the total binding energy (A), van der Walls contribution to the total MM-GBSA binding energy (B) and the electrostatic contribution to the total binding free energy (C). MM-GBSA contributions are evaluated using 1,000 samples from the simulations of P5S-1H1 complex with RBD, pdb id 7XS8. The residue-based decomposition for P5S-2B10/RBD binding shows the total binding energy (D), van der Walls contribution to the total MM-GBSA binding energy (E) and the electrostatic contribution to the total binding free energy (F). MM-GBSA contributions are evaluated using 1,000 samples from simulations of P5S-2B10 complex, pdb id 7XSC. Residue-based binding free energy values are shown for total energies as magenta-colored filled bars, van der Waals contributions as orange-colored bars and electrostatic contributions as light-brown colored bars.

Combined with a narrower spectrum of binding hotspots compared to BD55-1205, these data suggest that P52S-1H1 can be escaped by variants invoking mutations in positions K417, L455 and F486 (Figure 12A-C). Our results are consistent with the experiments of related class I IGHV3-53/3-66 antibodies binding to RBM and their shared escape mutant K417A/E/N/T mutations can lead to loss of neutralizing activity [110]. Moreover, the experimental single-site alanine scanning mutagenesis conducted in that study for the 15 epitope residues shared among IGHV3-53/3-66 antibodies showed that Y421A and F456A have broad impact on all related antibodies [110]. These experiments align well with the computational analysis revealing potential vulnerabilities and escape mutations for P5S-1H1. For P5S-2B10 the major binding hotspots are Y505, N487, Y489, R403, F486, A475, L455 and K417 (Figure 12D-F). The favorable van der Waals contacts are dominated by Y505, Y421, F456, A475 and F486 residues (Figure 12E), while electrostatics is driven by R403 and K417 (Figure 12F) While there are some notable differences in relative order of major energetic centers, the overall interaction pattern is similar reflecting strong dependence on conserved sites indispensable for stability and ACE2 binding as well as importance of L455, F456, A475, F486 and K417 (Figure 12D-F).

A recurring theme across all antibodies is the emergence of convergent evolutionary hotspots, particularly L455, F456 and also F486 which consistently serve as dominant escape hotspots. Functional constraints further underscore the importance of residues like Y421 and Y489, which are critical for RBD stability and ACE2 binding, making them less prone to mutational escape despite their importance in antibody binding. The results highlight trade-offs between epitope breadth, binding specificity, and adaptability to mutations. BD55-1205 with broad epitope coverage and distributed hotspot mechanisms, exhibit superior resilience to mutations. In contrast, BD-604 and OMI-42, with localized binding mechanisms, are more vulnerable to escape mutations at L455, F456 and A475 positions .P5S-1H1 and P5S-2B10 can be similarly vulnerable to mutations in these sites and additionally show sensitivity to a range of mutations in K417 (Figure 12). The shared vulnerabilities of P5S-1H1 and P5S-2B10 to mutations at L455 and F456, common across Class I antibodies, align with findings from experimental studies emphasizing convergent evolutionary pressures enabling robust escape [110]. The general insights obtained from MM-GBSA analysis and mutational scanning results highlight the role of convergent evolution, trade-offs between breadth and specificity in immune escape and functional stability constraints. L455 and F456 consistently emerge as dominant escape hotspots across all antibodies, enabling viral evolution through enhanced immune evasion. Antibodies with broad epitope coverage such as BD55-1205 exhibit greater resilience to mutations owing to a larger and more broadly distributed spectrum of major binding hotspots. We suggest that through this distributed hotspot mechanism BD55-1205 can sustain strong binding affinity and mitigate the effects of mutations in latest variants emerging in positions L455, F456 and A475.

## Discussion

Through an integrative approach combining structural analysis, mutational profiling, coarse-grained simulations, and MM-GBSA computations, we uncover critical insights into the binding epitopes, energetic contributions, and dynamic signatures of class I antibodies such as BD55-1205, BD-604, OMI-42, P5S-1H1, and P5S-2B10. These results not only elucidate the diversity of binding mechanisms but also underscore the nuanced interplay between antibody specificity, mutational adaptability, and viral evolution. The dynamic signatures of the RBD induced by different class I antibodies reveal important distinctions in their functional profiles. BD55-1205 induces significant stabilization of key RBD regions, effectively locking the RBD into a rigid conformation that enhances stability and strengthens binding interactions. This rigidity likely contributes to BD55-1205 superior ability to block ACE2 interactions and neutralize the virus. In contrast, BD-604 allows greater fluctuations in residues 450–475, suggesting a less constrained RBD conformation that may reduce its efficacy in blocking ACE2 binding. Intermediate behavior is observed for antibodies like P5S-1H1 and P5S-2B10, which stabilize the RBD to some extent but exhibit moderate fluctuations in regions like 490–505, indicating partial stabilization. These differences highlight the importance of targeting similar epitopes through distinct interaction mechanisms and patterns to counteract the virus ability to evolve resistance.

A central theme emerging from this work is the critical role of epitope breadth and interaction diversity in determining an antibody resilience to mutations. BD55-1205 antibody exemplifies the advantages of broad epitope coverage and distributed hotspot mechanisms. By engaging an extensive network of residues across the RBD, BD55-1205 minimizes its dependence on individual side-chain conformations, allowing it to maintain robust binding even when key residues are mutated. This adaptability is particularly evident in its tolerance to mutations at positions such as L455 and F456, which severely compromise other antibodies. The ability of BD55-1205 to sustain cumulative interactions underscores the importance of targeting diverse epitopes through multiple interaction mechanisms, a strategy that enhances resistance to immune evasion while maintaining functional integrity. The computational predictions generated through mutational scanning and MM-GBSA analysis demonstrate excellent agreement with experimental data on average antibody escape scores.

BD-604 demonstrates the vulnerabilities inherent in localized binding mechanisms. While BD-604 forms strong interactions with specific residues, its narrower focus renders it more susceptible to escape mutations at critical positions. For instance, mutations such as K417N, N460K, V445P, and F486S significantly reduce BD-604 efficacy against variants like BA.4/BA.5, BQ.1.1, and XBB. These findings highlight the trade-offs between specificity and adaptability in antibody design, emphasizing the need for strategies that balance high-affinity binding with resilience to antigenic variation. The weaker cumulative interactions observed for BD-604 also underscore the limitations of relying heavily on side-chain contacts, which are more sensitive to structural changes in the RBD. The shared vulnerabilities of P5S-1H1 and P5S-2B10 to mutations at L455 and F456 highlight their susceptibility to convergent evolutionary pressures. While they may retain efficacy against certain variants, their narrower spectrum of binding hotspots makes them less robust in the face of antigenic drift.

A recurring theme across all antibodies is the emergence of convergent evolutionary hotspots, particularly L455, F456, and F486, which consistently serve as dominant escape hotspots. These residues are indispensable for both ACE2 binding and antibody recognition, reflecting their dual role in mediating host-virus interactions and immune evasion. Functional constraints further underscore the importance of residues like Y421 and Y489, which are critical for RBD stability and ACE2 binding, making them less prone to mutational escape despite their importance in antibody binding. The study underscores the delicate balance between viral adaptation and immune evasion. The convergence of mutations in L455 and F456 highlights the intense selective pressures driving viral evolution, as the virus seeks to optimize immune evasion while maintaining fitness. Understanding these evolutionary constraints is essential for anticipating future mutations and designing therapeutics that target conserved residues and diverse epitopes. The subtle yet critical differences in epitope composition, residue dependence, structural dynamics, and mutational tolerance collectively shape the functional profiles of these antibodies, offering valuable insights for future research and therapeutic innovation.

## Conclusions

This study provides critical insights into the molecular mechanisms governing the binding of Class I neutralizing antibodies—BD55-1205, BD-604, OMI-42, P5S-1H1, and P5S-2B10—to the SARS-CoV-2 RBD. Despite targeting overlapping epitopes, these antibodies exhibit differences in their interaction networks, reliance on specific residues, and adaptability to mutations, collectively shaping their efficacy and resilience to viral evolution. BD55-1205 stands out for its broad epitope coverage and distributed hotspot mechanism, which enhance its ability to withstand mutations and maintain high-affinity binding. This resilience is attributed to its extensive backbone-mediated hydrogen bonds and hydrophobic interactions, enabling it to mitigate the impact of individual mutations. In contrast, BD-604 and OMI-42 demonstrate greater vulnerability to escape mutations due to their narrower focus and weaker cumulative interactions. P5S-1H1 and P5S-2B10 occupy an intermediate position, stabilizing the RBD to some extent but lacking the robustness of BD55-1205. The emergence of convergent mutations, particularly at residues L455 and F456, reflects the intense selective pressures driving viral adaptation. These positions serve as dominant escape hotspots, underscoring their dual role in mediating both host-virus interactions and immune recognition. These findings highlight the delicate interplay between structural flexibility, conserved residues, and evolutionary constraints, emphasizing the need for nuanced strategies to combat emerging variants. The subtle yet critical differences in epitope composition, residue dependence, and mutational tolerance collectively shape the functional profiles of these antibodies. Broad-spectrum antibodies like BD55-1205, which leverage extensive interaction networks and target conserved residues, represent valuable tools for addressing the ongoing threat posed by SARS-CoV-2. By integrating computational predictions with experimental validation, this study may provide basis for computational predictions of future mutations and escape pathways as well as support rational engineering of antibodies to broaden their epitope coverage or stabilize critical interactions which could enhance their resilience to prospective mutations.

**Supplementary Materials:** Figure S1 presents COVID-19 Variant Dashboard – Global coverage. Figure S2 presents overview of the phylogenetic analysis and divergence of Omicron variants. Figure S3 describes the evolutionary tree of current SARS-CoV-2 clades. Figure S4 presents representative conformations of ensembles of the RBD complexes and binding epitopes for class I antibodies obtained from the equilibrium trajectories. Figure S5 describes structural organization of the RBD complexes and binding epitopes for structures of BD-604 antibody with RBD and various Omicron variants. Figure S6 presents the results of dynamic mutational profiling of the RBD intermolecular interfaces in the different structures RBD complexes with BD-604. Figure S7 presents detailed results of the dynamic mutational profiling of the RBD intermolecular interfaces in the RBD complex with OMI-42. Table S1 lists Omicron variants with assigned clade annotation. Table S2 lists the intermolecular interfacial residues and contacts in the structure of BD55-1205 complex with RBD. Table S3 lists the intermolecular interfacial residues and contacts in the structure of BD-604 complex with RBD. Table S4 lists the intermolecular interfacial residues and contacts in the structure of OMI-42 complex with RBD. Table S5 lists the intermolecular interfacial residues and contacts in the structure of P5S-1H1 complex with RBD. Table S6 lists the intermolecular interfacial residues and contacts in the structure of P5S-2B10 complex with RBD.

## Author Contributions

Conceptualization, G.V.; Methodology, M.A., V.P., B.F., G.V.; Software, M.A., V.P., B.F., G.V.; Validation, G.V.; Formal analysis, G.V., M.A., V.P.,B.F.; Investigation, G.V. ; Resources, G.V., M.A., G.V. ; Data curation, M.A., G.C., G.V.; Writing— original draft preparation, G.V.; Writing—review and editing, G.V.; Visualization, .A.; G.V. Supervision G.V. Project administration, G.V.; Funding acquisition, G.V. All authors have read and agreed to the published version of the manuscript.

## Data Availability Statement

The original contributions presented in this study are included in the article/supplementary material. Crystal structures were obtained and downloaded from the Protein Data Bank (http://www.rcsb.org). The rendering of protein structures was done with interactive visualization program UCSF ChimeraX package (https://www.rbvi.ucsf.edu/chimerax/) and Pymol (https://pymol.org/2/) . All mutational heatmaps were produced using the developed software that is freely available at https://alshahrani.shinyapps.io/HeatMapViewerApp/.

## Acknowledgments

The authors acknowledge support from Schmid College of Science and Technology at Chapman University for providing computing resources at the Keck Center for Science and Engineering.

