## Supplemental Figures S1-S7, Tables S1-S6 for "Conformational Landscaping and Dynamic Mutational Profiling of Binding Interactions and Immune Escape for Broadly Neutralizing Class I Antibodies with SARS-CoV-2 Spike Protein: Distributed Binding Hotspot Networks Underlie Mechanism of Viral Resistance Against Existing Variants"

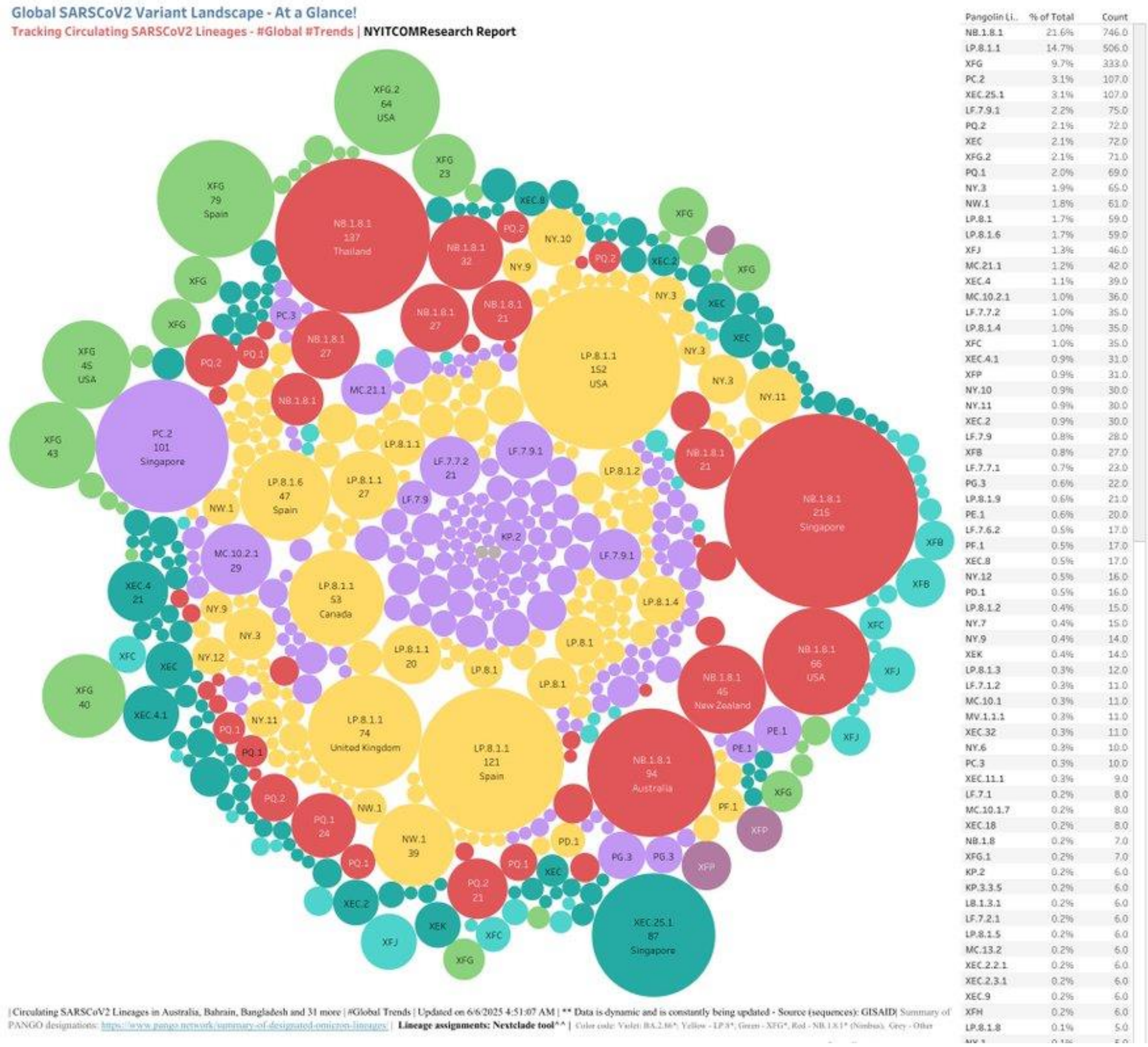


**Figure S1.** COVID-19 Variant Dashboard . Top lineages are NB.1.8.1 (21.6%), LP.8.1.1 (14.7%), XFG (9.7%), PC.2 (3.1%), XEC.25.1 (3.1%), LF.7.9.1 (2.2%), PQ.2 (2.1%), XEC (2.1%) XFG.2 (2.1%)(https://public.tableau.com/app/profile/raj.rajnarayanan/viz/USAVariantDB/VariantDashboard)

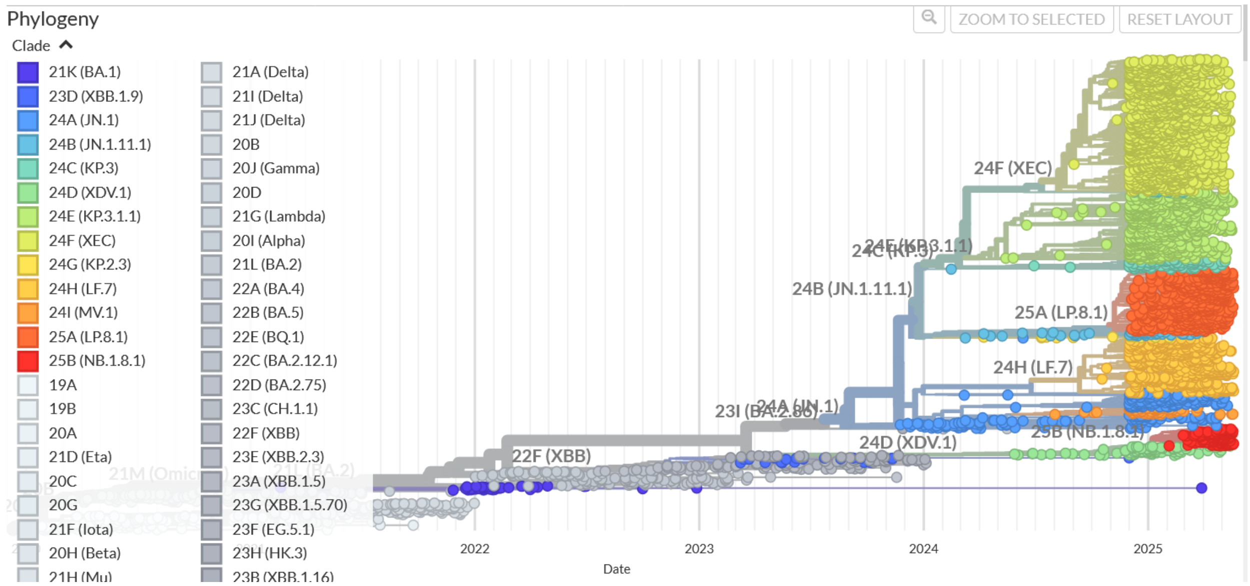


**Figure S2.** An overview of the phylogenetic analysis and divergence of Omicron variants. The graph is generated using Nextstrain, an open-source project for real time tracking of evolving pathogen populations (https://nextstrain.org/)

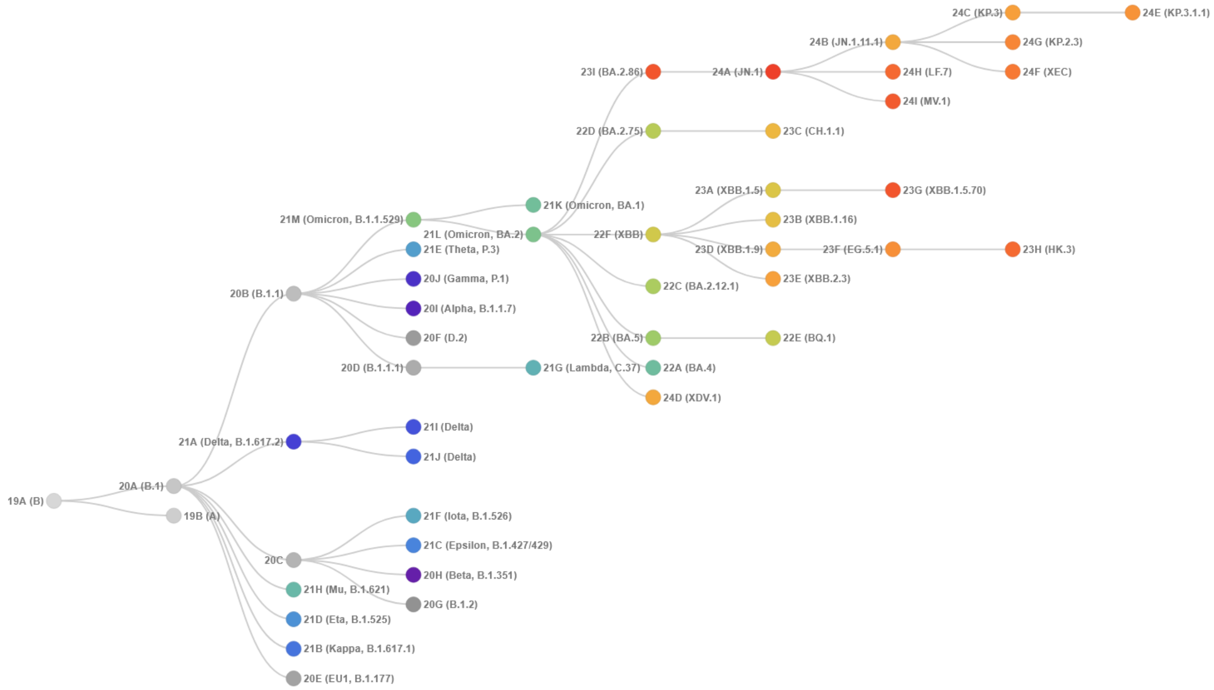


**Figure S3.** The evolutionary tree of current SARS-CoV-2 clades. The graph is generated using Nextstrain, an open-source project for real time tracking of evolving pathogen populations (https://nextstrain.org/). The clade 22F corresponds to XBB, 23A corresponds to XBB.1.5, 23G corresponds to XBB.1.5.70 (B.1.5+L455F+F456L) variant, 24C is KP.3 and 24E is KP.3.1.

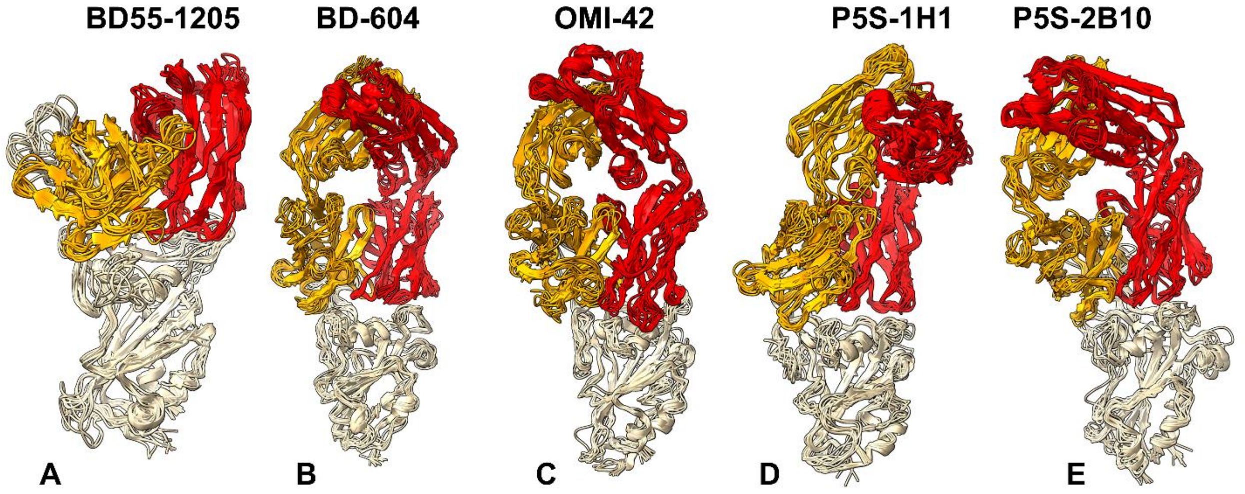


**Figure S4.** Conformational ensembles of the RBD complexes and binding epitopes for class I antibodies using representative conformations from the equilibrium trajectories. (A) The structure of BD55-1205 with XBB.1.5 RBD (pdb id 8XE9). The heavy chain in orange ribbons, the light chain in red ribbons. (B) The structure of BD-604 bound with BA.2 RBD (pdb id 8HWT). The heavy chain in orange ribbons, the light chain in red ribbons. (C) The structure of OMI-42 bound with Delta RBD (pdb id 8CBF). The heavy chain in orange ribbons, the light chain in red ribbons. (D) The structure of P5S-1H1 bound with RBD (pdb id 7XS8). The heavy chain in orange ribbons, the light chain in red ribbons. (IE) The structure of P5S-2B10 bound with RBD (pdb id 7XSC). The heavy chain in orange ribbons, the light chain in red ribbons.

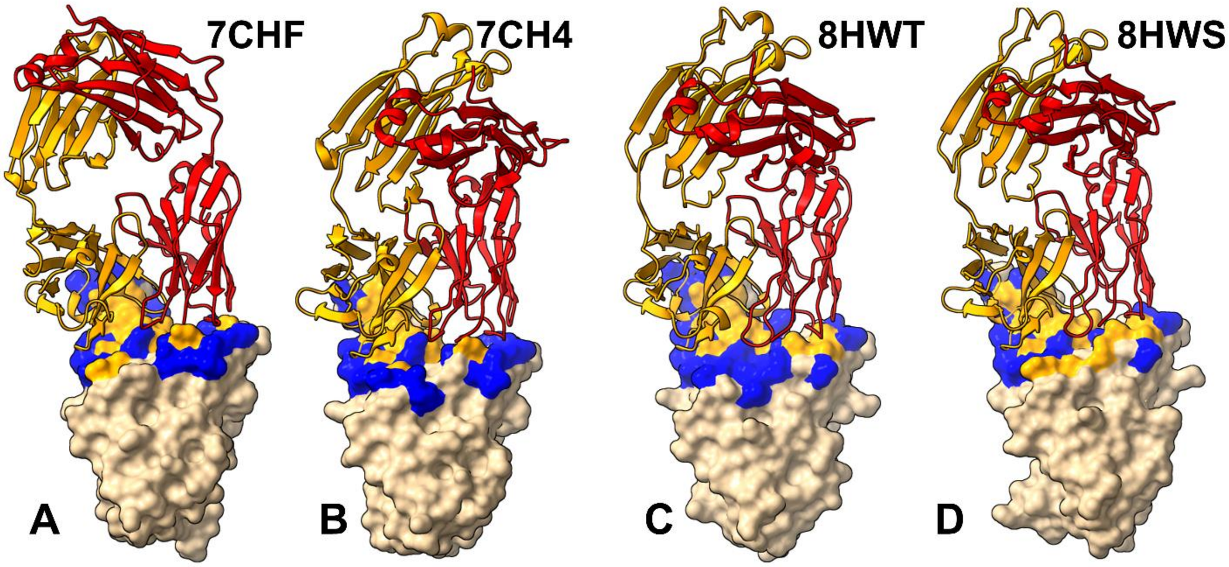


**Figure S5.** Structural organization of the RBD complexes and binding epitopes for structures of BD-604 antibody with RBD and various Omicron variants. The structures of BD-604 complex with RBD, pdb id 7CHF, (A) and pdb id 7CH4 (B), BD-604 complex with BA2 RBD, pdb id 8HWT (C) and BD-604 complex with BA.4/BA.5 RBD, pdb id 8HWS (D). The heavy chain in orange ribbons, the light chain in red ribbons. (D) The RBD and binding epitope footprint for BD-604. The binding epitope residues are shown in blue surface.

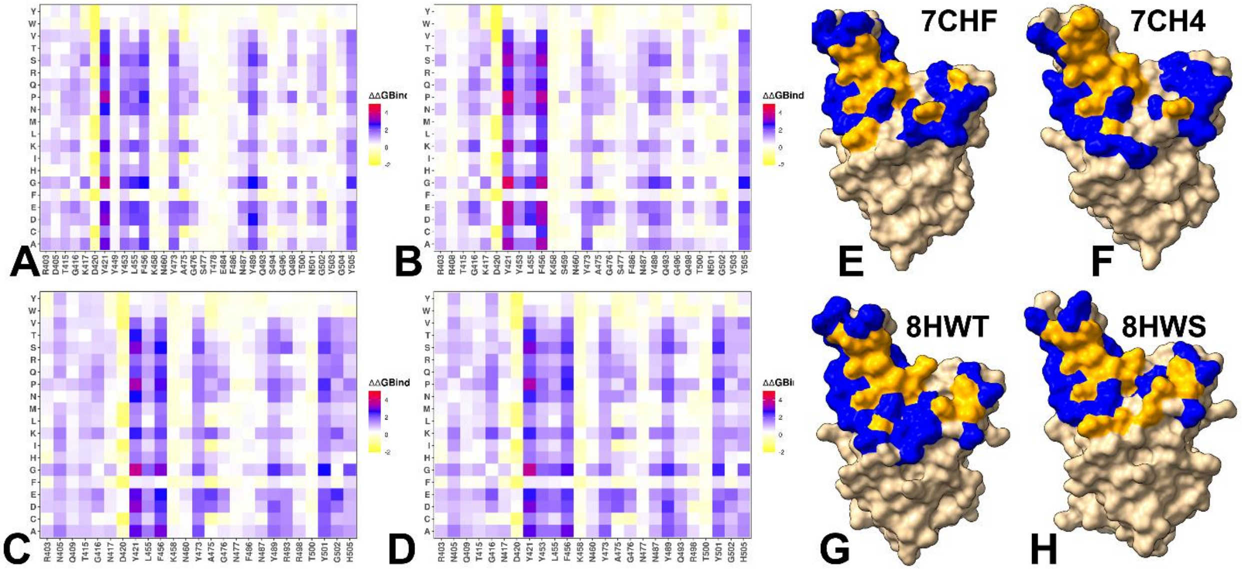


**Figure S6.** Ensemble-based dynamic mutational profiling of the RBD intermolecular interfaces in the different structures RBD complexes with BD-604. (A,B). The mutational scanning heatmaps are shown for the interfacial RBD residues in the BD-604 complex with RBD, pdb id 7CHF (A), BD-604 complex with RBD, pdb id 7CH4 (B), BD-604 complex with BA.2 RBD, pdb id 8HWT (C), BD-604 complex with BA.4/BA.5 RBD, pdb id 8HWS (D). The structures of BD-604 to RBD, pdb id 7CHF (E), pdb id 7CH4 (F), pdb id 8HWT (G), pdb id 8HWS (H). The heavy chain of BD-604 is in orange ribbons, and light chain is in red ribbons. The binding epitope is shown in blue surface and the positions of the RBD binding energy hotspots are shown in orange-colored surface.

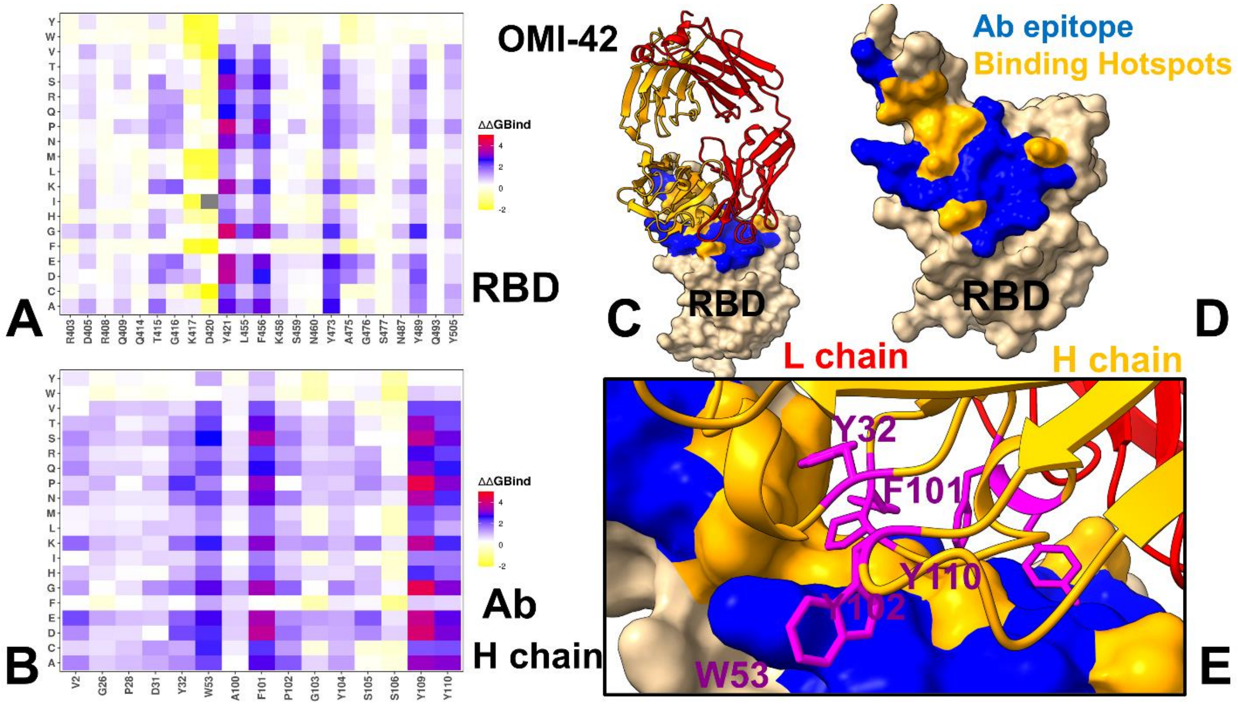


**Figure S7.** Ensemble-based dynamic mutational profiling of the RBD intermolecular interfaces in the RBD complex with OMI-42. (A,B). The mutational scanning heatmaps are shown for the interfacial RBD residues (A) and interfacial heavy chain residues of OMI-42 (B). The heatmaps show the computed binding free energy changes for 20 single mutations of the interfacial positions. (C) The structure of OMI-42 bound to RBD (pdb id 8CBF). The heavy chain of OMI-42 is in orange ribbons, and light chain is in red ribbons. The binding epitope is shown in blue surface and the positions of the RBD binding energy hotspots are shown in orange-colored surface. (D). RBD from the complex with OMI-42. The binding epitope residues are in blue surface and the binding interfacial RBD hotspots are in orange surface. (E) A closeup of the binding interface contacts of the OMI-42 hotspots from the heavy chain Y32, W53, F102 and Y110 . The heavy chain is in orange ribbons, light chain in red ribbons. The OMI-422 hotspots are shown in magenta sticks and annotated.

**Table S1. The list of Omicron Variants with Assigned Clade Annotation**

| clade | parent | Variant | WHO |
|---|---|---|---|
| 19B | 19A | 9A: 19A | |
| 20A | 19A | 19B: 19B | |
| 20B | 20A | 20A: 20A | |
| 20C | 20A | 20B: 20B | |
| 20D | 20B | 20C: 20C | |
| 20E | 20A | 20D: 20D | |
| 20F | 20B | 20E: 20E | |
| 20G | 20C | 20F: 20F | |
| 20H | 20C | 20G: 20G | Beta |
| 20I | 20B | 20I: 20I (Alpha) | Alpha |
| 20J | 20B | 20J: 20J (Gamma) | Gamma |
| 21A | 20A | 21A: 21A (Delta) | Delta |
| 21B | 20A | 21B: 21B (Kappa) | Kappa |
| 21C | 20C | 21C: 21C (Epsilon) | Epsilon |
| 21D | 20A | 21D: 21D (Eta) | Eta |
| 21E | 20B | 21E: 21E (Theta) | Theta |
| 21F | 20C | 21F: 21F (Iota) | Iota |
| 21G | 20D | 21G: 21G (Lambda) | Lambda |
| 21H | 20A | 21I: 21I (Delta) | Mu |
| 21I | 21A | 21H: 21H (Mu) | Delta |
| 21J | 21A | 21J: 21J (Delta) | Delta |
| 21K | 21M | 21K: 21K (BA.1) | Omicron |
| 21L | 21M | 21L: 21L (BA.2) | Omicron |
| 21M | 20B | 21M: 21M (Omicron) | Omicron |
| 22A | 21L | 22A: 22A (BA.4) | Omicron |
| 22B | 21L | 22B: 22B (BA.5) | Omicron |
| 22C | 21L | 22C: 22C (BA.2.12.1) | Omicron |
| 22D | 21L | 22D: 22D (BA.2.75) | Omicron |
| 22E | 22B | 22E: 22E (BQ.1) | Omicron |
| 22F | 21L | 22F: 22F (XBB) | Omicron |
| 23A | 22F | 23A: 23A (XBB.1.5) | Omicron |
| 23B | 22F | 23B: 23B (XBB.1.16) | Omicron |
| 23C | 22D | 23C: 23C (CH.1.1) | Omicron |
| 23D | 22F | 23D: 23D (XBB.1.9) | Omicron |
| 23E | 22F | 23E: 23E (XBB.2.3) | Omicron |
| 23F | 23D | 23F: 23F (EG.5.1) | Omicron |
| 23G | 23A | 23G: 23G (XBB.1.5.70) | Omicron |
| 23H | 23F | 23H: 23H (HK.3) | Omicron |
| 23I | 21L | 23I: 23I (BA.2.86) | Omicron |
| 24A | 23I | 24A: 24A (JN.1) | Omicron |
| 24B | 24A | 24B: 24B (JN.1.11.1) | Omicron |
| 24C | 24B | 24C: 24C (KP.3) | Omicron |

| | | | |
|---|---|---|---|
| 24D | 21L | 24D: 24D (XDV.1) | Omicron |
| 24E | 24C | 24E: 24E (KP.3.1.1) | Omicron |
| 24F | 24A | 24F: 24F (XEC) | Omicron |
| 24G | 24B | 24G: 24G (KP.2.3) | Omicron |
| 24H | 24A | 24H: 24H (LF.7) | Omicron |
| 24I | 24A | 24I: 24I (MV.1) | Omicron |
| 25A | 24B | 25A: 25A (LP.8.1) | Omicron |
| 25B | 24D | 25B: 25B (NB.1.8.1) | Omicron |

**Table S2.** The list of the intermolecular contacts in the structure of the BD55-1205 complex with RBD (pdb id 8XE9).

| RBD Residue | RBD Residue Number | RBD chain | Ab Residue | Ab Residue Number | Ab chain |
|---|---|---|---|---|---|
| ARG | 403 | C | ASN | 30 | B |
| ARG | 403 | C | GLY | 92 | B |
| ASN | 405 | C | ASP | 93 | B |
| THR | 415 | C | SER | 56 | A |
| THR | 415 | C | THR | 57 | A |
| THR | 415 | C | PHE | 58 | A |
| GLY | 416 | C | TYR | 52 | A |
| GLY | 416 | C | SER | 56 | A |
| GLY | 416 | C | PHE | 58 | A |
| ASN | 417 | C | TYR | 33 | A |
| ASN | 417 | C | TYR | 52 | A |
| ASN | 417 | C | TRP | 94 | B |
| ASN | 417 | C | PRO | 95 | B |
| ASP | 420 | C | TYR | 52 | A |
| ASP | 420 | C | SER | 56 | A |
| TYR | 421 | C | TYR | 33 | A |
| TYR | 421 | C | TYR | 52 | A |
| TYR | 421 | C | PRO | 53 | A |
| TYR | 421 | C | GLY | 54 | A |
| TYR | 421 | C | GLY | 55 | A |
| TYR | 453 | C | ILE | 101 | A |
| LEU | 455 | C | TYR | 33 | A |
| LEU | 455 | C | PRO | 53 | A |
| LEU | 455 | C | TRP | 94 | B |
| LEU | 455 | C | LEU | 99 | A |
| LEU | 455 | C | ILE | 101 | A |
| LEU | 455 | C | ARG | 102 | A |
| PHE | 456 | C | ARG | 31 | A |
| PHE | 456 | C | ASN | 32 | A |
| PHE | 456 | C | TYR | 33 | A |
| PHE | 456 | C | PRO | 53 | A |
| PHE | 456 | C | LEU | 99 | A |
| ARG | 457 | C | PRO | 53 | A |
| ARG | 457 | C | GLY | 54 | A |
| LYS | 458 | C | SER | 30 | A |
| LYS | 458 | C | ARG | 31 | A |
| LYS | 458 | C | PRO | 53 | A |
| LYS | 458 | C | GLY | 54 | A |
| SER | 459 | C | PRO | 53 | A |

| | | | | | |
|---|---|---|---|---|---|
| SER | 459 | C | GLY | 54 | A |
| LYS | 460 | C | GLY | 54 | A |
| LYS | 460 | C | GLY | 55 | A |
| LYS | 460 | C | SER | 56 | A |
| TYR | 473 | C | SER | 30 | A |
| TYR | 473 | C | ARG | 31 | A |
| TYR | 473 | C | ASN | 32 | A |
| TYR | 473 | C | PRO | 53 | A |
| GLN | 474 | C | ARG | 31 | A |
| ALA | 475 | C | PHE | 27 | A |
| ALA | 475 | C | THR | 28 | A |
| ALA | 475 | C | ARG | 31 | A |
| ALA | 475 | C | ASN | 32 | A |
| ALA | 475 | C | ARG | 97 | A |
| GLY | 476 | C | GLY | 26 | A |
| GLY | 476 | C | PHE | 27 | A |
| GLY | 476 | C | THR | 28 | A |
| GLY | 476 | C | ARG | 31 | A |
| GLY | 476 | C | ASN | 32 | A |
| ASN | 477 | C | GLY | 26 | A |
| ASN | 477 | C | PHE | 27 | A |
| ASN | 477 | C | THR | 28 | A |
| PRO | 486 | C | GLU | 104 | A |
| ASN | 487 | C | VAL | 2 | A |
| ASN | 487 | C | GLY | 26 | A |
| ASN | 487 | C | PHE | 27 | A |
| ASN | 487 | C | ARG | 97 | A |
| ASN | 487 | C | GLU | 104 | A |
| TYR | 489 | C | ASN | 32 | A |
| TYR | 489 | C | ARG | 97 | A |
| TYR | 489 | C | LEU | 99 | A |
| TYR | 489 | C | ARG | 102 | A |
| TYR | 489 | C | GLU | 104 | A |
| SER | 490 | C | ARG | 102 | A |
| PRO | 491 | C | ARG | 102 | A |
| LEU | 492 | C | ARG | 102 | A |
| GLN | 493 | C | ILE | 101 | A |
| GLN | 493 | C | ARG | 102 | A |
| ARG | 498 | C | SER | 31 | B |
| ARG | 498 | C | SER | 67 | B |
| THR | 500 | C | SER | 28 | B |
| THR | 500 | C | PHE | 29 | B |
| THR | 500 | C | GLY | 68 | B |

| | | | | | |
|---|---|---|---|---|---|
| TYR | 501 | C | SER | 28 | B |
| TYR | 501 | C | PHE | 29 | B |
| TYR | 501 | C | ASN | 30 | B |
| TYR | 501 | C | SER | 31 | B |
| GLY | 502 | C | SER | 28 | B |
| GLY | 502 | C | PHE | 29 | B |
| GLY | 502 | C | ASN | 30 | B |
| VAL | 503 | C | SER | 28 | B |
| HIS | 505 | C | SER | 28 | B |
| HIS | 505 | C | PHE | 29 | B |
| HIS | 505 | C | ASN | 30 | B |
| HIS | 505 | C | GLY | 92 | B |
| HIS | 505 | C | ASP | 93 | B |

**Table S3.** The list of the intermolecular contacts in the structure of the BD-604 complex with RBD (pdb id 8HWT).

| RBD Residue | RBD Residue Number | RBD chain | Ab Residue | Ab Residue Number | Ab chain |
|---|---|---|---|---|---|
| ARG | 403 | A | SER | 30 | L |
| ARG | 403 | A | ASP | 32 | L |
| ASN | 405 | A | ASN | 92 | L |
| ASN | 405 | A | SER | 93 | L |
| GLU | 406 | A | SER | 93 | L |
| GLN | 409 | A | SER | 93 | L |
| THR | 415 | A | SER | 56 | H |
| THR | 415 | A | PHE | 58 | H |
| GLY | 416 | A | TYR | 52 | H |
| GLY | 416 | A | SER | 56 | H |
| GLY | 416 | A | PHE | 58 | H |
| ASN | 417 | A | TYR | 33 | H |
| ASN | 417 | A | TYR | 52 | H |
| ASP | 420 | A | TYR | 52 | H |
| ASP | 420 | A | SER | 56 | H |
| TYR | 421 | A | TYR | 33 | H |
| TYR | 421 | A | TYR | 52 | H |
| TYR | 421 | A | SER | 53 | H |
| TYR | 421 | A | GLY | 54 | H |
| TYR | 453 | A | PRO | 101 | H |
| LEU | 455 | A | TYR | 33 | H |
| LEU | 455 | A | LEU | 99 | H |
| LEU | 455 | A | GLY | 100 | H |
| LEU | 455 | A | PRO | 101 | H |
| LEU | 455 | A | TYR | 102 | H |
| PHE | 456 | A | SER | 31 | H |
| PHE | 456 | A | TYR | 33 | H |
| PHE | 456 | A | ASP | 98 | H |
| PHE | 456 | A | LEU | 99 | H |
| PHE | 456 | A | TYR | 102 | H |
| ARG | 457 | A | SER | 53 | H |
| LYS | 458 | A | SER | 30 | H |
| LYS | 458 | A | SER | 31 | H |
| LYS | 458 | A | SER | 53 | H |
| LYS | 458 | A | GLY | 54 | H |
| SER | 459 | A | SER | 53 | H |
| SER | 459 | A | GLY | 54 | H |

| | | | | | |
|---|---|---|---|---|---|
| ASN | 460 | A | SER | 53 | H |
| ASN | 460 | A | GLY | 54 | H |
| ASN | 460 | A | GLY | 55 | H |
| ASN | 460 | A | SER | 56 | H |
| TYR | 473 | A | SER | 30 | H |
| TYR | 473 | A | SER | 31 | H |
| TYR | 473 | A | ASN | 32 | H |
| TYR | 473 | A | SER | 53 | H |
| GLN | 474 | A | SER | 31 | H |
| GLN | 474 | A | ASN | 32 | H |
| ALA | 475 | A | GLY | 26 | H |
| ALA | 475 | A | ILE | 27 | H |
| ALA | 475 | A | ILE | 28 | H |
| ALA | 475 | A | SER | 31 | H |
| ALA | 475 | A | ASN | 32 | H |
| ALA | 475 | A | ARG | 97 | H |
| GLY | 476 | A | GLY | 26 | H |
| GLY | 476 | A | ILE | 27 | H |
| GLY | 476 | A | ILE | 28 | H |
| GLY | 476 | A | SER | 31 | H |
| GLY | 476 | A | ASN | 32 | H |
| ASN | 477 | A | SER | 25 | H |
| ASN | 477 | A | GLY | 26 | H |
| ASN | 477 | A | ILE | 27 | H |
| ASN | 477 | A | ILE | 28 | H |
| LYS | 478 | A | GLY | 26 | H |
| PHE | 486 | A | VAL | 2 | H |
| PHE | 486 | A | ARG | 97 | H |
| PHE | 486 | A | ASP | 105 | H |
| PHE | 486 | A | VAL | 106 | H |
| ASN | 487 | A | VAL | 2 | H |
| ASN | 487 | A | GLY | 26 | H |
| ASN | 487 | A | ILE | 27 | H |
| ASN | 487 | A | ARG | 97 | H |
| TYR | 489 | A | ARG | 97 | H |
| TYR | 489 | A | LEU | 99 | H |
| TYR | 489 | A | TYR | 102 | H |
| TYR | 489 | A | ASP | 105 | H |
| PHE | 490 | A | TYR | 102 | H |
| LEU | 492 | A | TYR | 102 | H |
| ARG | 493 | A | SER | 31 | L |
| ARG | 493 | A | ASP | 32 | L |
| ARG | 493 | A | ALA | 50 | L |

| | | | | | |
|---|---|---|---|---|---|
| ARG | 493 | A | PRO | 101 | H |
| ARG | 493 | A | TYR | 102 | H |
| ARG | 498 | A | SER | 31 | L |
| ARG | 498 | A | SER | 67 | L |
| THR | 500 | A | SER | 67 | L |
| THR | 500 | A | GLY | 68 | L |
| TYR | 501 | A | GLY | 28 | L |
| TYR | 501 | A | SER | 30 | L |
| TYR | 501 | A | SER | 31 | L |
| TYR | 501 | A | SER | 67 | L |
| TYR | 501 | A | GLY | 68 | L |
| GLY | 502 | A | GLN | 27 | L |
| GLY | 502 | A | GLY | 28 | L |
| GLY | 502 | A | ILE | 29 | L |
| GLY | 502 | A | SER | 30 | L |
| VAL | 503 | A | GLN | 27 | L |
| HIS | 505 | A | GLY | 28 | L |
| HIS | 505 | A | ILE | 29 | L |
| HIS | 505 | A | SER | 30 | L |
| HIS | 505 | A | ASP | 32 | L |
| HIS | 505 | A | ASN | 92 | L |

**Table S4.** The list of the intermolecular contacts in the structure of the OMI-42 complex with RBD (pdb id 8CBF).

| RBD Residue | RBD Residue Number | RBD Chain | Ab Residue | Ab Residue Number | Ab Chain |
|---|---|---|---|---|---|
| ARG | 403 | E | GLU | 52 | L |
| ARG | 403 | E | LYS | 55 | L |
| ASP | 405 | E | ASN | 33 | L |
| GLU | 406 | E | TYR | 34 | L |
| ARG | 408 | E | GLY | 30 | L |
| ARG | 408 | E | GLY | 31 | L |
| GLN | 409 | E | TYR | 32 | L |
| GLN | 409 | E | TYR | 34 | L |
| GLN | 414 | E | TYR | 32 | L |
| THR | 415 | E | TYR | 32 | L |
| THR | 415 | E | TYR | 34 | L |
| THR | 415 | E | TYR | 93 | L |
| THR | 415 | E | GLY | 95 | L |
| THR | 415 | E | ASN | 96 | L |
| THR | 415 | E | TYR | 109 | H |
| GLY | 416 | E | TYR | 32 | L |
| GLY | 416 | E | TYR | 34 | L |
| GLY | 416 | E | TYR | 93 | L |
| GLY | 416 | E | TYR | 109 | H |
| LYS | 417 | E | TYR | 34 | L |
| LYS | 417 | E | GLU | 52 | L |
| LYS | 417 | E | LYS | 55 | L |
| LYS | 417 | E | TYR | 109 | H |
| LYS | 417 | E | TYR | 110 | H |
| ILE | 418 | E | TYR | 34 | L |
| ASP | 420 | E | TYR | 93 | L |
| ASP | 420 | E | SER | 105 | H |
| ASP | 420 | E | TYR | 109 | H |
| TYR | 421 | E | PRO | 102 | H |
| TYR | 421 | E | GLY | 103 | H |
| TYR | 421 | E | TYR | 104 | H |
| TYR | 421 | E | SER | 105 | H |
| TYR | 421 | E | SER | 106 | H |
| TYR | 421 | E | TYR | 109 | H |
| TYR | 421 | E | TYR | 110 | H |
| TYR | 453 | E | GLU | 52 | L |
| TYR | 453 | E | LYS | 55 | L |

| | | | | | |
|---|---|---|---|---|---|
| TYR | 453 | E | TYR | 110 | H |
| ARG | 454 | E | TYR | 110 | H |
| LEU | 455 | E | PHE | 101 | H |
| LEU | 455 | E | SER | 106 | H |
| LEU | 455 | E | TYR | 110 | H |
| PHE | 456 | E | PHE | 101 | H |
| PHE | 456 | E | PRO | 102 | H |
| PHE | 456 | E | SER | 106 | H |
| PHE | 456 | E | TYR | 110 | H |
| ARG | 457 | E | PRO | 102 | H |
| ARG | 457 | E | GLY | 103 | H |
| ARG | 457 | E | TYR | 104 | H |
| LYS | 458 | E | ASP | 30 | H |
| LYS | 458 | E | ASP | 31 | H |
| LYS | 458 | E | TRP | 53 | H |
| LYS | 458 | E | PRO | 102 | H |
| LYS | 458 | E | GLY | 103 | H |
| LYS | 458 | E | TYR | 104 | H |
| SER | 459 | E | TRP | 53 | H |
| ASN | 460 | E | TYR | 104 | H |
| TYR | 473 | E | PRO | 28 | H |
| TYR | 473 | E | ASP | 31 | H |
| TYR | 473 | E | TYR | 32 | H |
| TYR | 473 | E | PRO | 102 | H |
| GLN | 474 | E | PRO | 28 | H |
| ALA | 475 | E | VAL | 2 | H |
| ALA | 475 | E | GLY | 26 | H |
| ALA | 475 | E | PHE | 27 | H |
| ALA | 475 | E | PRO | 28 | H |
| ALA | 475 | E | TYR | 32 | H |
| ALA | 475 | E | LYS | 98 | H |
| ALA | 475 | E | ALA | 100 | H |
| GLY | 476 | E | VAL | 2 | H |
| GLY | 476 | E | GLY | 26 | H |
| GLY | 476 | E | PHE | 27 | H |
| GLY | 476 | E | PRO | 28 | H |
| SER | 477 | E | GLU | 1 | H |
| SER | 477 | E | GLY | 26 | H |
| SER | 477 | E | PHE | 27 | H |
| ASN | 487 | E | VAL | 2 | H |
| ASN | 487 | E | GLY | 26 | H |
| TYR | 489 | E | TYR | 32 | H |
| TYR | 489 | E | ALA | 100 | H |

| | | | | | |
|---|---|---|---|---|---|
| TYR | 489 | E | PHE | 101 | H |
| GLN | 493 | E | LYS | 55 | L |
| GLY | 504 | E | ASN | 33 | L |
| TYR | 505 | E | ASN | 33 | L |
| TYR | 505 | E | GLU | 52 | L |
| TYR | 505 | E | VAL | 53 | L |
| TYR | 505 | E | SER | 54 | L |

**Table S5.** The list of the intermolecular contacts in the structure of the P5S-1H1 complex with RBD (pdb id 7XS8).

| RBD Residue | RBD Residue Number | Residue Chain | Ab Residue | Ab Residue Number | Residue Chain |
|---|---|---|---|---|---|
| ARG | 403 | E | PHE | 32 | L |
| ASP | 405 | E | ASN | 92 | L |
| GLU | 406 | E | ASP | 93 | L |
| ARG | 408 | E | PHE | 58 | A |
| GLN | 409 | E | ASP | 93 | L |
| THR | 415 | E | PHE | 58 | A |
| THR | 415 | E | THR | 57 | A |
| THR | 415 | E | SER | 56 | A |
| GLY | 416 | E | TYR | 52 | A |
| GLY | 416 | E | SER | 56 | A |
| GLY | 416 | E | PHE | 58 | A |
| LYS | 417 | E | ASP | 93 | L |
| LYS | 417 | E | TYR | 33 | A |
| LYS | 417 | E | GLN | 100 | A |
| LYS | 417 | E | TYR | 52 | A |
| ASP | 420 | E | SER | 56 | A |
| ASP | 420 | E | PHE | 58 | A |
| ASP | 420 | E | TYR | 52 | A |
| TYR | 421 | E | TYR | 52 | A |
| TYR | 421 | E | SER | 53 | A |
| TYR | 421 | E | GLY | 55 | A |
| TYR | 421 | E | GLY | 54 | A |
| TYR | 421 | E | TYR | 33 | A |
| TYR | 421 | E | SER | 56 | A |
| TYR | 449 | E | ASN | 31 | L |
| TYR | 453 | E | VAL | 101 | A |
| TYR | 453 | E | PHE | 32 | L |
| ARG | 454 | E | TYR | 33 | A |
| LEU | 455 | E | GLN | 100 | A |
| LEU | 455 | E | VAL | 101 | A |
| LEU | 455 | E | TYR | 33 | A |
| LEU | 455 | E | LEU | 99 | A |
| PHE | 456 | E | TYR | 33 | A |
| PHE | 456 | E | TYR | 102 | A |
| PHE | 456 | E | LEU | 99 | A |

| | | | | | |
|---|---|---|---|---|---|
| PHE | 456 | E | ASP | 98 | A |
| PHE | 456 | E | GLN | 100 | A |
| PHE | 456 | E | ASN | 32 | A |
| ARG | 457 | E | SER | 53 | A |
| ARG | 457 | E | GLY | 54 | A |
| LYS | 458 | E | SER | 31 | A |
| LYS | 458 | E | GLY | 54 | A |
| LYS | 458 | E | SER | 30 | A |
| LYS | 458 | E | SER | 53 | A |
| SER | 459 | E | GLY | 54 | A |
| SER | 459 | E | SER | 53 | A |
| ASN | 460 | E | GLY | 54 | A |
| ASN | 460 | E | SER | 56 | A |
| ASN | 460 | E | GLY | 55 | A |
| ASN | 460 | E | SER | 53 | A |
| TYR | 473 | E | ASN | 32 | A |
| TYR | 473 | E | SER | 30 | A |
| TYR | 473 | E | SER | 53 | A |
| TYR | 473 | E | SER | 31 | A |
| GLN | 474 | E | SER | 31 | A |
| ALA | 475 | E | SER | 31 | A |
| ALA | 475 | E | GLY | 26 | A |
| ALA | 475 | E | ILE | 27 | A |
| ALA | 475 | E | THR | 28 | A |
| ALA | 475 | E | ASN | 32 | A |
| ALA | 475 | E | ARG | 97 | A |
| GLY | 476 | E | THR | 28 | A |
| GLY | 476 | E | ASN | 32 | A |
| GLY | 476 | E | SER | 31 | A |
| GLY | 476 | E | GLY | 26 | A |
| GLY | 476 | E | ILE | 27 | A |
| SER | 477 | E | GLY | 26 | A |
| SER | 477 | E | THR | 28 | A |
| GLU | 484 | E | TYR | 102 | A |
| PHE | 486 | E | GLY | 26 | A |
| PHE | 486 | E | ASP | 105 | A |
| PHE | 486 | E | VAL | 106 | A |
| PHE | 486 | E | ARG | 97 | A |
| PHE | 486 | E | ILE | 27 | A |
| PHE | 486 | E | VAL | 2 | A |
| ASN | 487 | E | GLY | 26 | A |
| ASN | 487 | E | LEU | 99 | A |
| ASN | 487 | E | ILE | 27 | A |

| | | | | | |
|---|---|---|---|---|---|
| ASN | 487 | E | ASP | 105 | A |
| ASN | 487 | E | THR | 28 | A |
| ASN | 487 | E | ASN | 32 | A |
| ASN | 487 | E | ARG | 97 | A |
| TYR | 489 | E | ARG | 97 | A |
| TYR | 489 | E | TYR | 102 | A |
| TYR | 489 | E | LEU | 99 | A |
| TYR | 489 | E | ASN | 32 | A |
| TYR | 489 | E | ASP | 105 | A |
| PHE | 490 | E | TYR | 102 | A |
| GLN | 493 | E | VAL | 101 | A |
| GLN | 493 | E | TYR | 102 | A |
| SER | 494 | E | PHE | 32 | L |
| TYR | 495 | E | PHE | 32 | L |
| TYR | 495 | E | SER | 30 | L |
| GLY | 496 | E | PHE | 32 | L |
| GLY | 496 | E | SER | 30 | L |
| PHE | 497 | E | SER | 30 | L |
| GLN | 498 | E | SER | 30 | L |
| GLN | 498 | E | GLY | 68 | L |
| GLN | 498 | E | SER | 67 | L |
| GLN | 498 | E | ASN | 31 | L |
| THR | 500 | E | SER | 67 | L |
| THR | 500 | E | THR | 69 | L |
| THR | 500 | E | ILE | 29 | L |
| THR | 500 | E | GLN | 27 | L |
| THR | 500 | E | GLY | 28 | L |
| THR | 500 | E | GLY | 68 | L |
| ASN | 501 | E | GLY | 68 | L |
| ASN | 501 | E | ILE | 29 | L |
| ASN | 501 | E | SER | 30 | L |
| ASN | 501 | E | GLY | 28 | L |
| GLY | 502 | E | GLN | 27 | L |
| GLY | 502 | E | GLY | 28 | L |
| GLY | 502 | E | ILE | 29 | L |
| VAL | 503 | E | GLN | 27 | L |
| GLY | 504 | E | GLN | 27 | L |
| TYR | 505 | E | HIS | 90 | L |
| TYR | 505 | E | ILE | 2 | L |
| TYR | 505 | E | ASN | 92 | L |
| TYR | 505 | E | GLN | 27 | L |
| TYR | 505 | E | GLY | 28 | L |
| TYR | 505 | E | ILE | 29 | L |

**Table S6.** The list of the intermolecular contacts in the structure of the P5S-2B10 complex with RBD (pdb id 7XSC).

| RBD Residue | RBD Residue Number | RBD Chain | Ab Residue | AB Residue Number | AB Chain |
|---|---|---|---|---|---|
| ARG | 403 | E | ASN | 93 | B |
| ARG | 403 | E | ASP | 92 | B |
| ASP | 405 | E | GLN | 27 | B |
| ASP | 405 | E | ASN | 93 | B |
| THR | 415 | E | THR | 57 | A |
| THR | 415 | E | SER | 56 | A |
| THR | 415 | E | PHE | 58 | A |
| GLY | 416 | E | TYR | 52 | A |
| GLY | 416 | E | PHE | 58 | A |
| GLY | 416 | E | SER | 56 | A |
| LYS | 417 | E | ASP | 92 | B |
| LYS | 417 | E | TYR | 33 | A |
| LYS | 417 | E | TYR | 52 | A |
| LYS | 417 | E | TYR | 100 | A |
| ASP | 420 | E | PHE | 58 | A |
| ASP | 420 | E | TYR | 52 | A |
| ASP | 420 | E | SER | 56 | A |
| TYR | 421 | E | SER | 56 | A |
| TYR | 421 | E | SER | 53 | A |
| TYR | 421 | E | TYR | 33 | A |
| TYR | 421 | E | TYR | 52 | A |
| TYR | 421 | E | GLY | 54 | A |
| TYR | 421 | E | GLY | 55 | A |
| TYR | 453 | E | ASP | 92 | B |
| TYR | 453 | E | PHE | 32 | B |
| LEU | 455 | E | SER | 53 | A |
| LEU | 455 | E | TYR | 33 | A |
| LEU | 455 | E | TYR | 100 | A |
| PHE | 456 | E | SER | 53 | A |
| PHE | 456 | E | TYR | 33 | A |
| PHE | 456 | E | SER | 31 | A |
| PHE | 456 | E | TYR | 100 | A |
| PHE | 456 | E | ASN | 32 | A |
| PHE | 456 | E | GLY | 101 | A |
| ARG | 457 | E | SER | 53 | A |
| ARG | 457 | E | GLY | 54 | A |
| LYS | 458 | E | ARG | 71 | A |

| | | | | | |
|---|---|---|---|---|---|
| LYS | 458 | E | SER | 31 | A |
| LYS | 458 | E | SER | 53 | A |
| LYS | 458 | E | SER | 30 | A |
| LYS | 458 | E | GLY | 54 | A |
| LYS | 458 | E | GLY | 55 | A |
| SER | 459 | E | GLY | 54 | A |
| SER | 459 | E | SER | 53 | A |
| ASN | 460 | E | SER | 56 | A |
| ASN | 460 | E | GLY | 54 | A |
| ASN | 460 | E | GLY | 55 | A |
| TYR | 473 | E | ASN | 32 | A |
| TYR | 473 | E | SER | 31 | A |
| TYR | 473 | E | SER | 53 | A |
| TYR | 473 | E | SER | 30 | A |
| GLN | 474 | E | SER | 31 | A |
| ALA | 475 | E | PHE | 27 | A |
| ALA | 475 | E | ARG | 97 | A |
| ALA | 475 | E | THR | 28 | A |
| ALA | 475 | E | ASN | 32 | A |
| ALA | 475 | E | SER | 31 | A |
| GLY | 476 | E | ASN | 32 | A |
| GLY | 476 | E | GLY | 26 | A |
| GLY | 476 | E | PHE | 27 | A |
| GLY | 476 | E | THR | 28 | A |
| SER | 477 | E | THR | 28 | A |
| SER | 477 | E | PHE | 27 | A |
| GLY | 485 | E | ASP | 102 | A |
| PHE | 486 | E | VAL | 2 | A |
| PHE | 486 | E | GLY | 26 | A |
| PHE | 486 | E | ASP | 102 | A |
| PHE | 486 | E | ARG | 97 | A |
| ASN | 487 | E | ASN | 32 | A |
| ASN | 487 | E | THR | 28 | A |
| ASN | 487 | E | GLY | 26 | A |
| ASN | 487 | E | ASP | 102 | A |
| ASN | 487 | E | PHE | 27 | A |
| ASN | 487 | E | ARG | 97 | A |
| CYS | 488 | E | ASP | 102 | A |
| TYR | 489 | E | ASN | 32 | A |
| TYR | 489 | E | ASP | 102 | A |
| TYR | 489 | E | ARG | 97 | A |
| TYR | 489 | E | GLY | 101 | A |
| GLY | 496 | E | ARG | 30 | B |

| | | | | | |
|---|---|---|---|---|---|
| GLN | 498 | E | ARG | 30 | B |
| THR | 500 | E | ARG | 30 | B |
| THR | 500 | E | ASP | 28 | B |
| ASN | 501 | E | ARG | 30 | B |
| ASN | 501 | E | ASP | 28 | B |
| GLY | 502 | E | ARG | 30 | B |
| GLY | 502 | E | ASP | 28 | B |
| TYR | 505 | E | ASN | 93 | B |
| TYR | 505 | E | PHE | 32 | B |
| TYR | 505 | E | ILE | 29 | B |
| TYR | 505 | E | ARG | 30 | B |
| TYR | 505 | E | ASP | 28 | B |
| TYR | 505 | E | ASP | 92 | B |
